# Sharing Massive Biomedical Data at Magnitudes Lower Bandwidth Using Implicit Neural Function

**DOI:** 10.1101/2022.12.03.518948

**Authors:** Runzhao Yang, Tingxiong Xiao, Yuxiao Cheng, Anan Li, Jinyuan Qu, Rui Liang, Shengda Bao, Xiaofeng Wang, Jue Wang, Jinli Suo, Qingming Luo, Qionghai Dai

**Author notes:** **Materials and Correspondence** Correspondence and material requests should be addressed to Jinli Suo. these authors contributed equally to this work (Runzhao Yang, Tingxiong Xiao, Yuxiao Cheng, Anan Li).

## Abstract

Efficient storage and sharing of massive biomedical data would open up their wide accessibility to different institutions and disciplines. However, compressors tailored for natural photos/videos are rapidly limited for biomedical data, while emerging deep learning based methods demand huge training data and are difficult to generalize. Here we propose to conduct Biomedical data compRession with Implicit nEural Function (BRIEF) by representing the original data with compact neural networks, which are data specific and thus have no generalization issues. Benefiting from the strong representation capability of implicit neural function, BRIEF achieves 2 ∼ 3 orders of magnitude compression on diverse biomedical data at significantly higher fidelity than existing techniques. Besides, BRIEF is of consistent performance across the whole data volume, supports customized spatially-varying fidelity. BRIEF’s multi-fold advantageous features also serve reliable downstream tasks at low bandwidth. Our approach will facilitate low-bandwidth data sharing, and promote collaboration and progress in the biomedical field.

## Introduction

The wide accessibility of biomedical data is fundamental for the flourishing of modern biomedical science^1–3^. On the one hand, advanced optical imaging technologies^4–10^ and emerging computational imaging schemes^11–15^ are producing a massive and rapidly growing amount of high-throughput, high-quality data. On the other hand, it is now very common for researchers with an informatics background to develop new algorithms and even benchmark new technologies. Researchers have devised numerous informatics-based techniques^16–19^ for the image processing and data mining, aiming to extract, compare, search, and effectively manage vast biomedical datasets^20^. A number of tools, systems and resources have been specifically produced for handling very large image datasets^21,22^ as well. Efficient data sharing of vast biomedical data would assist interdisciplinary teamwork to accelerate biomedical discoveries via data reanalysis. Confronted with massive data, researchers have to use low-resolution acquisitions or even selectively keep the important ones due to storage limitation^23^. Besides, the data sharing via either local hard disk or network is burdensome because of insufficient storage space or transmission bandwidth. Some large data-sharing sites use servers with high bandwidth and large storage space^24,25^, which alleviates the sharing challenges on the server side at the expense of high maintenance costs but is incapable of solving the downloading bandwidth and storage problems on the user side. These issues bring significant costs and greatly hamper the progress of biomedical studies.

The explosive biomedical data demands high-performance data compression to fit the available storage/bandwidth. After decades of development, some commercial compression tools can achieve high fidelity representation at high compression ratios^26^, but these widely used compressors such as H.264^27^ and H.265^28^ are tailored for natural scenes and cannot readily compress biomedical data, which is of largely different characteristics from natural scenes and would suffer from either low compression ratio or severe artifacts that would harm the downstream tasks. To compress data other than natural photos or videos, some deep learning based approaches are gaining momentum^29–31^ and achieve superior performance to conventional techniques, benefiting from the supervised model training that learns the proper compression specifications from a large amount of data. In spite of the high performance, current deep learning based compression is limited in multiple aspects: (i) The compressor is tailored for the training data and cannot serve as a general compression handling diverse biomedical data. Even for the same data type, slight variations in imaging settings necessitate retraining a new model, which involves time-consuming training data acquisition and model training. (ii) The model is tuned towards fixed compression quality, so one can not flexibly trade-off between compression ratio and fidelity without retraining. Even with several models at different compression ratios, one can only choose among several discrete levels, while a flexible control at fine levels or even continuously is apparently a better option. (iii) In many biomedical studies only part of the data is worthy of high-fidelity storage/transmission or useful for downstream tasks, but the highly desired spatially-varying compression quality, either defined manually or automatically, is not supported.

Implicit neural function (INF)^32^ uses a neural network to parameterize a continuous signal, such as shape^33–35^ and scene^36–38^, by mapping spatial or spatio-temporal coordinate towards the corresponding data intensity. Compared to conventional representations defined over a discrete grid, implicit neural function is able to model fine details beyond the grid resolution^32,39^. Since most digital biomedical data intrinsically describes a continuous signal/field, implicit neural function holds great potential to find the low dimensional latent structure underlying the original data and achieve high fidelity data compression^40–47^, i.e., fit the data with a small number of network parameters but at high representation accuracy. Moreover, such fitting is conducted in data specific manner and is thus generally applicable for diverse data, without the generalization issue confronted by previous deep learning based data compression techniques. To sum up, in comparison to conventional methods and state-of-the-art deep learning approaches, INF offers a continuous parameterization that potentially leads to higher compression ratios. Moreover, INF circumvents the generalization issue and enhances flexibility, thus establishing the rationale for employing INF to compress massive biomedical data.

In this article, we propose a general and flexible biomedical data compression approach built on implicit neural function—BRIEF, utilizing implicit neural function’s powerful representation capability for compact data encoding. To cover diverse biomedical data in a unified manner, we design two adaptive strategies—adaptive data partitioning decomposing the data volume into blocks with similar complexity, and adaptive parameter allocation according to the features of different blocks. Implemented under an optimization framework, BRIEF searches for the best neural representation under constraints from the target file size and pre-defined importance map, and thus supports precise specification of the compression ratio and saliency-guided compression quality. Experimentally, BRIEF reaches 2~3 orders of magnitude compression ratio with largely superior performance to existing compression tools on diverse biomedical data, such as neurons, clinical CT data, X-ray holographic nano-tomographic data, and MRI, etc. Moreover, most network hyperparameters in BRIEF are consistent across diverse data, which facilitates easy-to-use for non-expert users. The proposed BRIEF and its features largely alleviate the urgent need for sharing and storage of enormous amounts of high-quality biomedical data, and would contribute to the entire community of biomedical research.

## Results

### The principle of implicit neural function for compression and decompression

The schematic of BRIEF is shown in Fig. 1(a). For a large digital biomedical data volume, we regard it as a discrete sampling of a continuous signal defined over a spatial domain and attempt to find a function with a small number of parameters to describe the mapping from the coordinate to the corresponding intensity, i.e., to serve as a compact encoding of the volumetric data. Considering its strong expression power, here we use implicit neural function as the mapping function to achieve powerful, flexible, and general data compression for biomedical data.

**Fig. 1.**
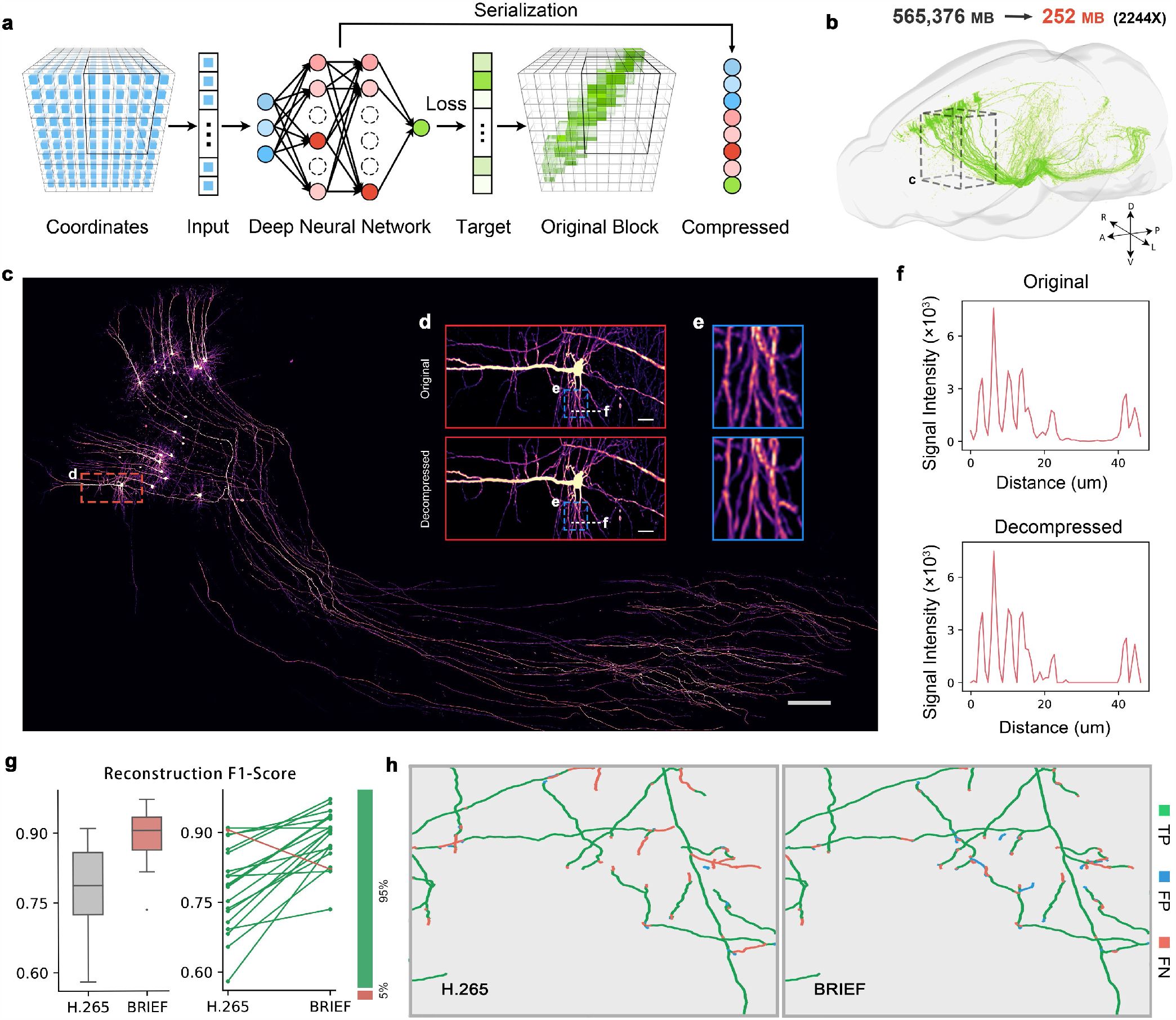
The schematic of BRIEF and its compression performance on the whole-mouse-brain neuronal population. **a**, During BRIEF compression, the huge target data volume is firstly adaptively partitioned into blocks with similar complexity (details of partitioning strategies described in Supplementary Figure 1), and undergo separate compression via pursuing a neural network that maps the voxel coordinates to their intensity values. The pursued best matching network is serialized into a bit sequence for low bandwidth transmission or storage. **b**, The fluorescently labeled neurons across a whole mouse brain, including 28965 ×20005 ×12000 voxels with 16 bit intensity values. BRIEF can achieve around 2244 ×compression (from 565,376 to 252MB) at high data fidelity. **c**, Zoomed-in view of the compressed region-of-interest (ROI) labeled in **b**. Scale bar: 200*μm*. **d**,**e**, Zoomed-in view of the highlighted regions in **c** and **d** respectively, in parallel with the MIP of the original data. Scale bar: 20*μm*. **f**, Intensity profile of the original data and its decompressed counterpart along the white dashed line labeled in **d. g**, The mean and standard variance of Reconstruction F1-score on the traced 3D morphological structure after compression by H.265 and BRIEF, and the performance difference over 20 samples, in which the BRIEF performs better and worse are highlighted with green and red lines respectively. **h**, The error map of traced 3D nerve fibers on the decompressed data. TP (True Positive): the correctly reconstructed structures; FP (False Positive): the incorrectly hallucinated structures; FN (False Negative): the missing details.

Although a neural network can fit an arbitrarily complex function with an infinite number of parameters, statistically strong correlations exist only locally, so it is more strategic/efficient to decompose a complex function into simple ones and use a series of small neural networks to fit these frequency-concentrated components^48,49^. Therefore, we adaptively divide the original data into blocks and pursue such a mapping function within each local region, as shown in Supplementary Figure 1(a). For each block, BRIEF first generates a coordinate system matching the original data volume, e.g., a uniform grid (2D, 3D, or even 4D) shown in Fig. 1(a) where each blue particle represents a grid point. Then, BRIEF builds a multi-layer perceptron (MLP) as a parameterized mapping function defined on the grid to fit the original data, with the number and numerical precision of network parameters in the MLP determining the size of the compressed data. Naturally, data compression is formulated as an optimization problem, where the weights and biases of the neurons in the neural network are optimized to match the network’s output towards the intensity of the original biomedical data across the grid. The intensity is either scalar (single-channel data) or vector (multi-channel data), and here we show a single-channel example for presentation clarity. The optimization can be conducted using either batch gradient descent or mini-batch gradient descent, depending on the available memory size. After finalizing the neural network, the network parameters (including the weights and biases of each neuron) and side information (including the hyperparameters of the neural network structure, the spatial range of the data block, bit depth, etc.) are extracted to form the compressed representation by serialization. Finally, the compressed data of different blocks are concatenated for successive storage or transmission. Experimentally, the compression ratio can achieve up to 2 ∼ 3 magnitudes at high fidelity for most biomedical data. The data decompression can be conducted simply by applying forward propagation to the spatial coordinates by corresponding implicit neural function, parsed from the compressed representation and side information. Afterward, the decompression of different blocks is conducted separately, and the reconstructed blocks are stitched together to form the whole data volume, as shown in Supplementary Figure 1(b).

As for the block-wised compression of the large data volume, building instance-specific partitioning and storage/bandwidth allocation among blocks is necessary to cover the large diversity among varying biomedical data. To address this issue we propose a two-stage adaptive strategy: Firstly, an adaptive partitioning method is designed to decompose the original data into a proper number of blocks, of different sizes and similar complexities (Extended Data Fig. 2). Followed by this, BRIEF allocates parameters according to the characteristics of each data block, and dynamically adjusts the parameter allocation to balance the compression quality over all the blocks during the compression of each data block (Extended Data Fig. 3).

Benefiting from the novel coding paradigm leveraging high representation power of implicit neural function and adaptive divide-and-conquer strategy, BRIEF not only achieves compression capability beyond existing compression tools, but also possesses more distinguished features, which enable high generalization ability, wide applicability, stability, flexibility, and scalability:

- BRIEF pursues the optimal neural representation tailored for the target data, and is thus totally free of generalization issues confronted by existing deep learning based compressors.
- BRIEF is a general compression approach widely applicable for diverse biomedical data for both biologic studies and clinical diagnosis, such as 2D images, and 3D volume.
- BRIEF produces stable compression quality along different dimensions since the compression network is directly defined over the homogeneous spatial grid to model the anatomical structure of the target data.
- Spatially varying compression fidelity can be tailored according to the region’s importance, which remains a key challenge for conventional compression methods.
- The compression ratio can be precisely regulated by pursuing a proper neural encoding (i.e., network structure, weights, and bias of a set of MLPs) under strict budget constraints from available storage or transmission bandwidth.
- Our partitioning strategy not only enhances BRIEF’s representation accuracy but also makes BRIEF scalable for the size of the original data and allows parallel computation, which enhances computation efficiency.

### BRIEF compresses whole-mouse-brain neural population by 3 magnitudes with high data fidelity

A neuroanatomical study of the brain-wide neural architecture can provide rich insights into the intrinsic structure and working mechanism of the brain’s neural system. However, the huge number of neurons and high dynamic range causes massive data volume, which hampers the subsequent analysis and demands a compressor with high fidelity at a high compression ratio. We apply BRIEF to the whole-mousebrain fluorescent neural data captured by the HD-fMOST system^9^, with 0.32×0.32 ×1.00*μm*^3^ resolution to resolve the elaborate neural structures and 16-bit recording to cover the 5-magnitudes fluorescent intensity. The final 3D volume is 28965 ×20005× 12000 voxels and occupies 565,376 MegaBytes (MB), as shown in Fig. 1(b).

BRIEF can achieve 2244× compression (from 565,3769MB to 252MB) while preserving the complex thin structures and high-dynamic-range fluorescent intensities faithfully, as shown in Fig. 1(c), the zoomed-in view showing the maximum intensity projection (MIP) of a region-of-interest (ROI) labeled in Fig. 1(b). For a better demonstration of the compression capability, we display the further zoomed-in view of an ROI in Fig. 1(c) side by side with the original version, as shown in Fig. 1(d) and (e). From the comparison, one can see that BRIEF preserves the complex and faint morphological structures of whole-mouse-brain neurons with high precision despite such a high compression ratio. Besides the visual comparison, we also plot in Fig. 1(f) the profiles of fluorescence signal across a highlighted line with large intensity variations to validate BRIEF’s high fidelity. We also demonstrate our performance on the huge whole-brain neuron data in Supplementary Video 1.

Thanks to the high-fidelity compression of BRIEF, researchers can conduct subsequent downstream processing after low-band-width data sharing. For neurons, the single most important task is to precisely trace the morphological structure of the nerve fibers and soma. At a high compression ratio up to 2244, our method still preserves the spatial location of nerve fibers. We used the open-source software Gtree^1^ to locate the nerve fibers with 3D tracing. Compared with H.265 we not only reconstruct the nerve fibers correctly (more TP) and ensure their continuity (less FN), but also avoid the wrong hallucination (less FP), as shown in Fig. 1(h). For numerical comparison, we use the F1-score to measure the preserved morphological structures in the 3D tracing results after compression. Our algorithm significantly improves the accuracy of the 3D traces and outperforms H.265 in 95% cases (Fig. 1(g)).

### BRIEF achieves effective compression on diverse clinical computed tomography (CT) data

The continuously and rapidly growing medical image data size is putting high pressure on hospitals’ storage capacity and hampering data sharing for intelligent medical data analysis. These clinical data consist of rich details crucial for diagnosis and demand high-quality compression.

To demonstrate BRIEF’s performance on clinical imaging data, we used the HiP-CT^10^ dataset, a collection of CT data of human organs. The HiP-CT dataset consists of 54 three-dimensional data volumes from different human organs or organ parts. Such clinical CT data are from tissue to cellular scales, and are of large diversity in appearance, with some typical examples shown in the top show of Fig. 2(a).

**Fig. 2.**
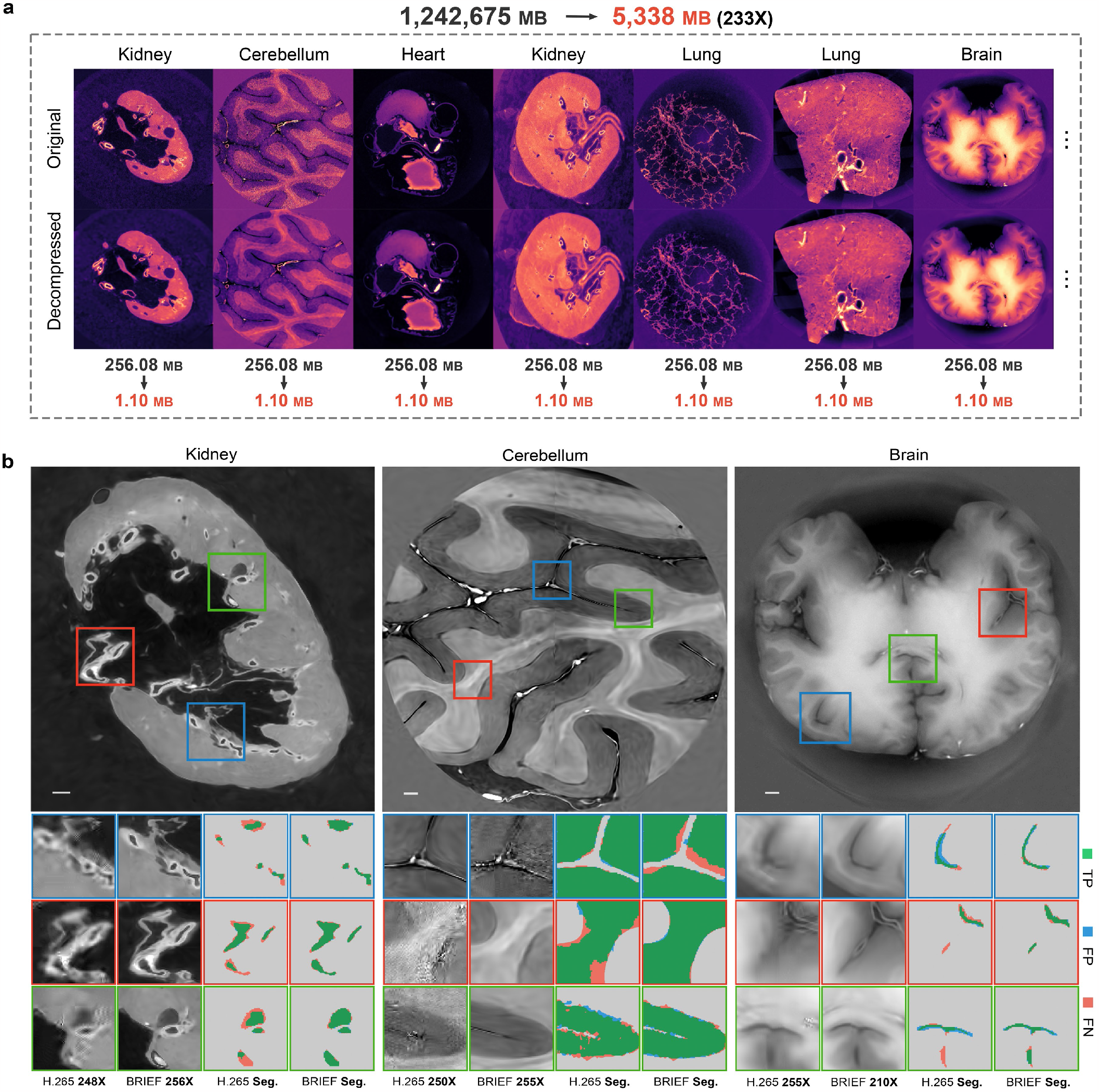
The compression capability of BRIEF on organic CT data from HiP-CT dataset and the performance of downstream tasks. **a**, Exemplar 2D slices from seven data groups of different organs or organ parts (cropped into 512 ×512 ×512 voxels) and their decompressed counterparts after compression by BRIEF (bottom). The whole data collection is compressed by 233 × (from 1,242,675MB to 5,338MB), and the compression ratio for different groups are labeled below the exemplar images. **b**, The comparison of segmentation results on decompressed data by BRIEF and H.265 for kidney, cerebellum, and brain, at around 230 ×compression ratio. For each example, the zoomed-in view of decompressed data and segmentation error map of highlighted regions are shown side by side, with segmentation results color-coded by TP, FP, and FN respectively. Scale bars from left to right: 4*mm*, 2*mm*, 5*mm*.

The whole dataset occupies 1,242,675 MB in total, and after compression, the data size decreases to 5,338 MB (i.e., 233 × compression). From the comparison of the exemplar data in Fig. 2(a) top and their decompressed version in Fig. 2(a) bottom, one can see that BRIEF preserves the sharp edges and complex structures at high precision, such as the tubules in kidney, white and grey matters in cerebellum, bronchioli in lung, even under around 230 × compression ratios. One can zoom in or visit Supplementary Videos 2 and 3 for a clearer comparison.

For clinical data, the compression algorithm should neither destroy the key cues important for diagnosis (e.g., fine structure and sharp edges) nor produce any artifacts that might result in misdiagnosis. At a compression ratio around 230 ×, BRIEF reproduces the kidney’s structure including the sharp boundaries and widely distributed tiny tubules (Fig. 2(b), left), the white and grey matters in cerebellum (Fig. 2(b), middle) and the sulci in brain (Fig. 2(b), right) as well. We also evaluate the compression quality in terms of segmentation accuracy that helps diagnosis, mainly the boundaries between kidney tubules, between white and grey matter, and between cortex. Here we used marching squares^50^ and intelligent scissors^51^ to semi-automatically segment the ROIs, and the segmentation results are shown in the bottom row in Fig. 2(b). We also compare the segmentation accuracy with state-of-the-art compressor H.265. Clearly, after compression by H.265 the segmentation results suffer from large area false segmentation, especially near the boundaries. The incorrect segmentation is mainly attributed to the severe streak artifacts in the high-frequency regions, as shown in the decompressed data placed on the side. In contrast, BRIEF preserved the details with high fidelity and few regions are misclassified.

### BRIEF allows higher-fidelity compression for more important regions

For a biomedical data volume, the details at different locations are not equally important, e.g., the precision instrument operator would preserve the faintest observations, the researchers would focus on the regions that might reveal new insights, the doctors on those help diagnosis, etc. Such requirements demand a compression approach that supports customized spatially varying fidelity across the volume, while the existing compressor lacks flexibility for region-specific quality. For BRIEF, we can simply generate a weight map from user-defined ROIs to represent the priority and incorporate it into the optimization. It should be noted that specifying an ROI is not mandatory but optional in BRIEF.

For example, for the nerve fibers with faint intensities, we generate a binary weight map to prioritize such neurons (intensity within the range [100, 2000]), and conduct a comparison on tens of data volumes to show the advantages of introducing prioritized fidelity. The results are shown in Fig. 3(a)-(e), with “w Weight Map” denoting the result with weight map and “w/o Weight Map” denoting that sets equal importance. Using F1 with a threshold of 200 and Reconstruction F1-score as the evaluation metric, the box plots and ladder plots in Fig. 3(a) show that we can get both higher accuracy, more correct 3D tracing, and smaller variance as well in almost all the cases. One typical example is shown in Fig. 3(b), from which one can see that after imposing higher priority BRIEF preserves more faint signals (top row), and thus facilitates more accurate morphology reconstruction (bottom row). We also zoom in one highlighted subregion for clearer comparison in terms of MIP (Fig. 3(c)) and intensity profile along the dashed line (Fig. 3(d)). The results clearly show that BRIEF can be tuned for higher-fidelity compression for fainter signals and thus retaining more details around smaller nerve fibers, which consists well with the statistics in Fig. 3(a).

**Fig. 3.**
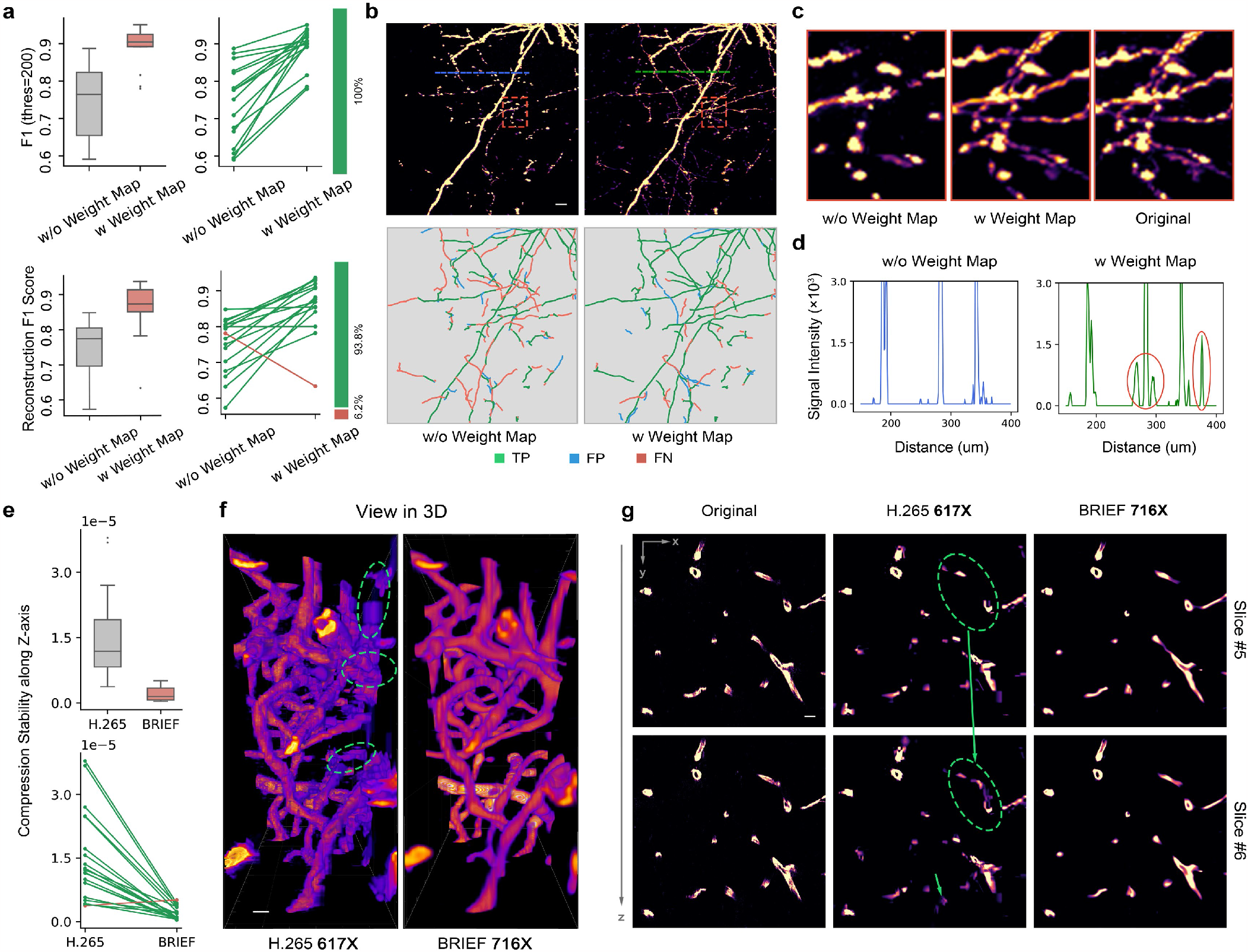
BRIEF’s features in supporting spatially varying quality specification and homogeneous compression quality along different dimensions. **a**, The mean and standard variance of the F1 with a threshold of 200 (top) and Reconstruction F1-score on the traced neuronal morphology (bottom), calculated from 16 data volumes decompressed by BRIEF with and without weight map. Here the BRIEF with weight map performs better and worse are highlighted with green and red lines respectively. **b**, The visual comparison of MIP (top) and tracing result (bottom) on one exemplar decompressed data in **a**, with the color-coded error map computed with respect to the morphology retrieved from the original data. Scale bar: 30*μm*. **c**, Zoomed-in view of the highlighted ROIs in **b**, in parallel with the original data. **d**, Intensity profiles along the dashed lines labeled in **b. e**, The mean and standard deviation of BRIEF’s and H.265’s performance fluctuations along axial slices, calculated from 20 test samples and displayed with the same layout and legend as in **a. f**, 3D vascular structures reconstructed from one of decompressed data in **e** by H.265 (left) and BRIEF (right), shown in *xz*-top view. Green dashed circles highlight the incorrectly broken and merged vessels caused by H.265’s unstable compression quality. Scale bar: 9*μm*. **g**, The visual comparison among original data, BRIEF’s and H.265’s decompressed version, on two adjacent slices from **f**, with compression ratio and slice index labeled at top and right respectively. The green dashed circle highlights H.265’s flicker artifacts along *z*-axis. Scale bar: 9*μm*.

It also potentially adds another advantageous feature to our approach, namely, users could specify the compression quality of ROIs according to the downstream tasks. Equipped with the ability to customize compression ratio together with ROI, BRIEF can make the most extensive use of the limited storage or transmission budget.

### BRIEF produces stable compression quality along each dimension/axis

For biomedical data that describes a volume in 3D space (e.g. neurons and vessels), it is important for the compression algorithm to keep stable compression quality along each axis to facilitate accurate reconstruction of its 3D structure. We compare the fluctuation range of compression quality along *z*-axis for vessels between BRIEF and H.265, with the results shown in Fig. 3(e)-(g). Compared to H.265, BRIEF has a much smaller performance variation along *z*-axis, and the superiority is consistent over almost all the test data. As highlighted by the green arrows, one can see that H.265 loses some details in slice #5 while hallucinates some in slice #6, which causes abrupt changes between adjacent slices in the decompressed data. Such artifacts will harm the successive 3D reconstruction of vascular structure and cause incorrect merging or even breaking of vessels. The artifacts by H.265 are mainly caused by flaws in optical flow prediction, which partially explains the performance limitation of H.265 for 3D biomedical data, which hinders further processing and subsequent diagnosis. Oppositely, BRIEF obtains results almost identical to the original input. It is mainly attributed to the fact that we use an implicit neural function to model the 3D structure embedded in the data and avoid these flicker effects.

### BRIEF supports precise specification of compression ratio

For effective management of massive biomedical data, it is crucially important to conduct compression to fit the large volumes into available storage or transmission bandwidth. An approach allowing users to set compression ratios freely can be helpful for making extensive use of the given budget. Supplementary Figure 2 demonstrates the ability of BRIEF to precisely control the compression ratio, which is largely different from five methods whose final achieved compression ratio might largely deviate from the target setting. In comparison, for each method, we give fixed parameter settings related to compression ratio (please refer to Supplementary Table 1 for details) and apply compression to the data randomly selected from the three aforementioned datasets. Obviously, for existing methods, even tuned for the same intended compression ratio, the achieved compression ratio varies among different data and the among-data variance is especially large for SGA and H.264. This will lead to the following shortcomings: First, repeated tuning is inevitable for users to approximate the expected compression ratio for the input data. Second, the final achieved ratio cannot exactly match the target ratio. Even worse, the compression setting reaching the expected compression ratio for data is inapplicable for others, and data-specific tuning is required. Oppositely, BRIEF returns a compression representation of the input data with the exact specified compression ratio (Extended Data Fig. 4). This virtue facilitates users choosing a compression model according to the available transmission bandwidth or storage. Besides, we can also search for a proper compressor to achieve a given quality by searching within a range of compression ratios.

### BRIEF achieves high-fidelity compression for a diverse of biomedical data

In addition to the neurons and clinical CT data described above, we have validated BRIEF’s wide applicability on diverse biomedical data, such as fluorescence micro-optical sectioning tomography of brain vessels^52^, X-ray holographic nano-tomography (XNH)^8^, MRI^53^, Fundus image^54^, bright-field microscopy^55^, two-photon microscopy^56^, and confocal microscopy^57^ data.

The vessel data were obtained by the HD-fMOST system^9^, with different fluorescent labeling from neurons. To precisely reconstruct 3D vascular structure from the decompressed data, the compression methods need to accurately preserve the 3D vascular structure, including the location, continuity, and thickness of the vessels. The reconstructed 3D vascular structure is displayed in Supplementary Figure 3 (right), which indicates that BRIEF can successfully preserve the intricate spatial vascular structures and enormous branches with uneven thicknesses under high compression ratio. On the contrary, some thin capillary vessels are lost after compression by H.265. Moreover, BRIEF exhibits a superior noise suppression effect, where noise comes from the capture of vessel data, indicating its high suppression of background noise while preserving a great amount of target signal (see Supplementary Note 3 for more experiments).

The XNH and MRI data were released from two published articles^8,53^ respectively. Similar to the CT data, these data also include rich details describing the fine structures crucial for successive medical diagnosis. BRIEF can achieve 2 magnitudes compression on these data while preserving fine structures, as shown in Supplementary Figures 5 and 6 and Supplementary Video To validate the generalization of BRIEF, we further test BRIEF’s compression quality on three other brain imaging modalities: bright-field microscopy^55^, two-photon microscopy^56^, and confocal microscopy^57^. The compression results are shown in Supplementary Figure 7. As a general compressor, BRIEF can theoretically compress visual information other than 3D volumetric data. For 2D image, we compare BRIEF with JPEG on Fundus^54^ data and show that BRIEF clearly preserves the continuous blood vessels and avoids color distortion, as demonstrated in Supplementary Figure 8.

### BRIEF outperforms the state-of-the-art (SOTA) compressors on biomedical data

To demonstrate the advantageous performance of our approach, here we conduct comparison on three groups of biomedical data (neurons, vessels, and organic CT) with six methods with state-of-the-art compression quality: four widely used commercial compression techniques—JPEG^58^, JPEG 2000^59^, H.264^27^, H.265^28^ and two representative machine learning schemes in the literature—SGA^29^ and DVC^60^. For quantitative performance comparison, we adopt three evaluation metrics for the decompressed data: F1 with thresholds for neurons and vessels, and PSNR and SSIM for organic CT. The performance statistics of six methods at different BPVs (bits per voxel of compressed data) are illustrated in Fig. 4(a). Generally, BRIEF shows superior performance to the SOTAs over a wide range of BPVs, across three groups of data and in terms of all the evaluation metrics. The superiority is more distinct at higher compression ratios (lower BPV). Moreover, compared with SOTAs, the deterioration of the data fidelity along with the growth of compression ratio is less observable in BRIEF, which surprisingly indicates less contradiction between BPV and quality metrics in our method. The implicit neural function, coupled with the proposed adaptive partitioning and parameter allocation mechanism, utilized in BRIEF demonstrates a notable advantage in mitigating fidelity loss as the BPV decreases. The inherent compatibility of implicit neural function with the continuous nature of biomedical data signals enhances its efficacy in capturing intricate details. Moreover, the proposed scheme conducts partitioning by maximizing the within-block correlation, thus enabling adaptive allocation of parameters based on the structure of the target data.

**Fig. 4.**
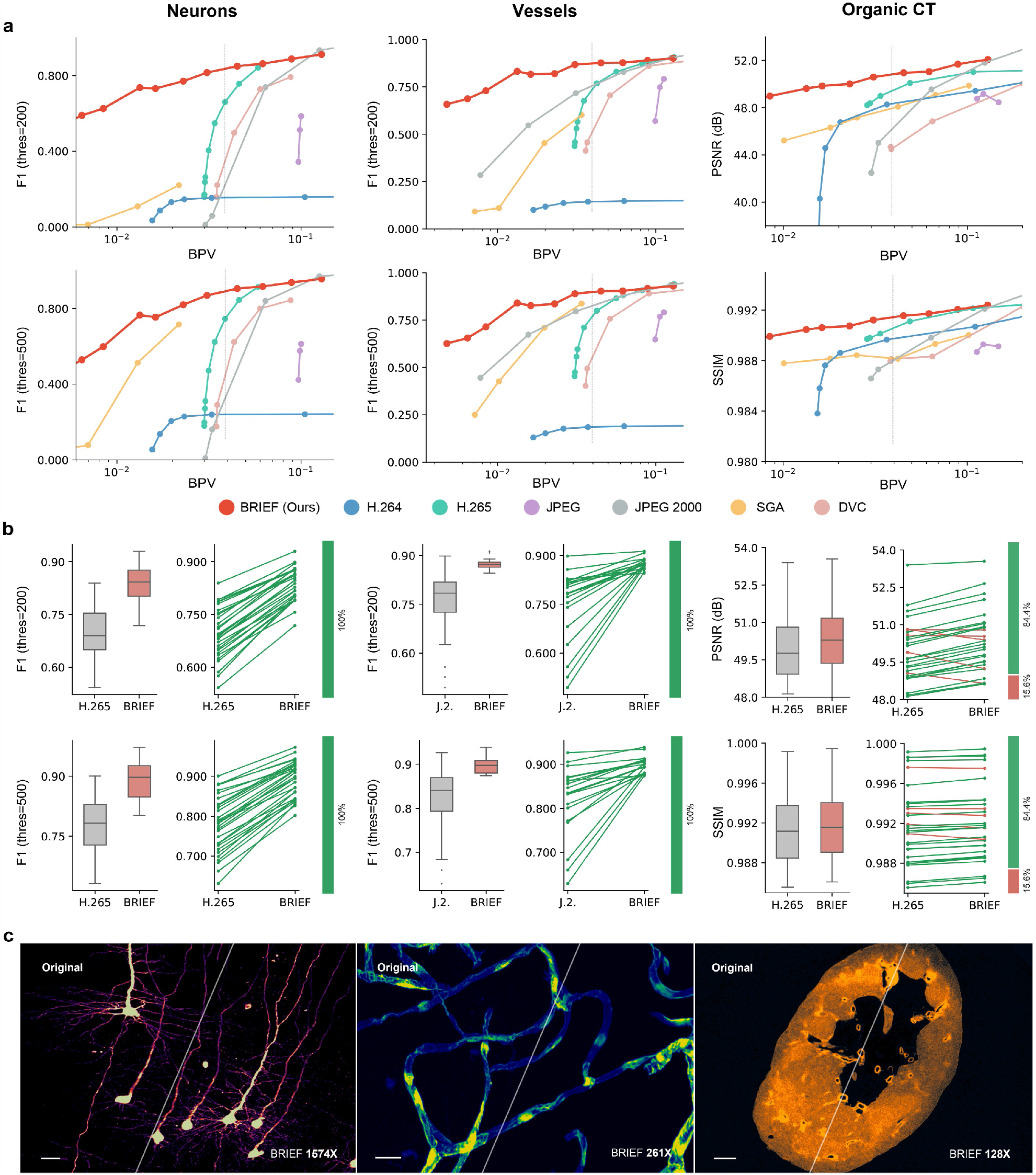
Performance comparison between BRIEF and state-of-the-art compressors. **a**, The performance of BRIEF and six SOTAs across three groups of biomedical data at varying compression ratios. The SOTAs include four widely used commercial compression tools (H.264, H.265, JPEG, JPEG 2000) and two emerging machine learning based compressors (SGA and DVC). For quantitative analysis, we use F1 with thresholds of 200 and 500 as evaluation metrics for neurons and vessels with sparse structures, while PSNR and SSIM for organic CT including rich details. We control the BPV within range [10^−2^, 10^−1^] and test at multiple discrete BPVs. **b**, The comparison between BRIEF and the second most competitor, around a given BPV annotated by grey vertical lines in **a**. Left: the mean and standard variance of compression quality over twenty test data. Right: the performance difference on each instance, with the ones BRIEF performs better and worse are respectively highlighted with green and red lines. **c**, The visual comparison of the original data and BRIEF’s decompressed version, with compression ratio labeled at the right bottom. Scale bars from left to right: 100*μm*, 12*μm*, 5*mm*.

To show more specific details in the comparison results, we compare with the second best method (H.265) on three different data (neurons, vessels, and organic CT) around specific BPVs, and the results are demonstrated in Fig. 4(b). In general, the average performance of BRIEF is higher than the second-best competitor, with a close or less deviation in all the cases. Specifically, BRIEF outperforms H.265 in (i) 100% cases in neurons, in terms of F1 with thresholds of 200 and 500, (ii) 100% cases in vessels, in terms of F1 with thresholds of 200 and 500, (iii) 84.4% cases in organic CT, considering the metric of PSNR and SSIM. The results not only prove the superior performance of BRIEF in most cases but a better generalization ability on different groups of biomedical data. Notably, we have compared the compression quality of BRIEF and SOTAs on the axonal fibers, which is presented in Supplementary Table 2 and Supplementary Figure 9.

## Discussion

We propose BRIEF, an efficient and easy-to-use compression approach for biomedical data compression based on neural network, to facilitate efficient storage of massive biomedical data, and timely sharing across disciplinary and institutional borders. BRIEF outperforms widely used compression methods which are tailored for natural photos or videos. The superior performance of BRIEF primarily arises from the rigorous mathematical derivation and proof of INF’s frequency concentration property in this manuscript, along with a tailored adaptive partitioning and allocation scheme built upon this property.

BRIEF intrinsically searches for a compact neural network representation fitting the original data under the available budget, and is also advantageous over deep learning-based compression in that it achieves higher-fidelity compression, requiring no training data and totally out of generalization problems.

In addition, BRIEF is built on an implicit neural function defined over the homogeneous spatial grid and thus achieves comparable fidelity along both lateral and axial dimensions. One can also specify a saliency map to tailor the fidelity across the volume, which is especially useful for the huge data with only limited regions of interest, or spatially varying contributions to the downstream tasks. Implemented in an optimization manner, BRIEF supports trade-off between fidelity and at arbitrary compression ratios instead of roughly choosing across coarse levels as in off-the-shelf compression software/tools.

Our approach is widely applicable and of marked performance for diverse biomedical data. We test BRIEF’s performance on ten types of biomedical data (fluorescence image of brain neurons and vessels, CT data of human organs, XNH data of drosophila leg, knee MRI, fundus image, calcium imaging of neural activities, bright-field microscopy, two-photon microscopy, and confocal microscopy data) and achieve 2∼ 3 magnitudes compression while keeping at high fidelity sufficient for downstream studies. The universality holds the potential for easy sharing of a large amount of valuable data in the whole community.

To give full play of intrinsic neural representation in data compression and cover large diversity in biomedical data, BRIEF proposed adaptive partitioning and adaptive budget (number of parameters) allocation. Such strategies enable high fidelity at high compression ratios and also allows parallel compression, which can be helpful due to the wide availability of multi-core processor and the maturity of multi-thread programming.

The total compression and decompression time for whole datasets are shown in Supplementary Table 3. The compression process of BRIEF need to solve optimization problems to fit the data under constraints from the given budget, including searching for the best partitioning strategy and progressive adjustment of the network parameters. Hence, it is slower than conventional ready tools (Supplementary Table 4 and Supplementary Figure 10). To address this issue, we can try to find better initialization or compress blocks in parallel by implementing the whole neural network using fully-fused CUDA kernels^61^. However, the decompression speed of BRIEF is almost one-magnitude faster than H.265 since we conduct decompression by simple forward propagation through the neural network, which is rapid and scalable. Although terabyte-scale data sets may scale up their decompression time, BRIEF’s is still much faster than state-of-the-art compressors (Supplementary Table 4 and Supplementary Figure 10). It is worth noting that biomedical data need to be compressed only once before saving or transmission but decompressed frequently for processing and analysis. Therefore, BRIEF is more efficient when used in practice in the biomedical field.

Our blocking strategy divides a large problem into smaller ones, which can then be solved with parallel computation. In our future work, we intend to further refine and optimize the allocation of computational resources, aiming to achieve fast compression. As future work, we are trying to integrate dynamic causal models to strengthen the interpretability of BRIEF and extract semantic information in the compression domain.

BRIEF contributes in multiple aspects: 1) opening the new applications of the emerging AI technique—INF, by aligning well its advantages with the demanding compression ratios and adaptability in biomedical data compression; 2) uncovering INF’s frequency concentration property, providing a theoretical guidance for data partitioning when handling large-volume data; 3) enhancing INF’s encoding capability, via proposing the adaptive data partitioning and parameter allocation scheme, for high compression quality.

To facilitate its easy use over the whole community, we have distributed the proposed compressor as an open-source software package with a detailed manual on the settings. We believe that it will provide the researchers with a powerful tool for data sharing, which would allow reliable verification of experimental results and innovative research built on the existing huge amount of biomedical data.

## Methods

### Network structure of this paper

In comparison to other multi-layer perceptron (MLP) architectures, the SIREN^32^-like MLP exhibits superior representation accuracy and more stable performance (Supplementary Figure 11). Consequently, in this paper we choose the SIREN-like MLP architecture. We use a SIREN-like MLP, consisting of an input layer, several hidden layers and an output layer, with the network structure shown in Extended Data Fig. 1. The input layer embeds the spatial coordinates of the original data to a higher dimensional space using a sinusoidal function with learnable parameters (frequency and phase). The sinusoidal function helps map the coordinates to the sine wave frequency components of biomedical data, which tend to lie on low-dimensional structures. The hidden layers compose a fully connected neural network with activation functions to model the intrinsic structure of data in the higher dimensional embedded space. While the output layer, a weighted linear combination with bias, i.e., without activation function, fits the mapping from the embedded coordinates to the final intensity. One can flexibly tune the number of hidden layers, activation function and the number of neurons in these layers. In our implementation, the network adopts the sinusoidal activation function and includes 5 hidden layers with equal number of neurons. The number of neurons is determined by the compression ratio and the size of the original data (Extended Data Fig. 4).

Denoting **v** as the normalized coordinate vector (2D, 3D, or even 4D) and *L* as the number of hidden layers, the output of the input layer **z**^(0)^, of the *l*th hidden layer **z**^(*l*)^, and the final output *f*_*θ*_ (**v**) are respectively defined as

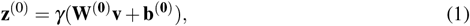

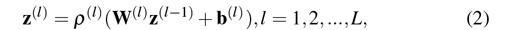

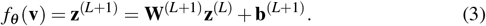

Here *γ*(·) is the sinusoidal mapping function; **W**^(*l*)^ is the weight, **b**^(*l*)^ is the bias, and *ρ*^(*l*)^(·) is the activation function; *f*_*θ*_ (**v**) is the final mapping from coordinate **v** to the corresponding intensity, with *θ* = {**W**^(*l*)^, **b**^(*l*)^ | *l* = 0, 1, …, *L* + 1} representing the learnable parameters of neural network. In the implementation, we adopted the sinusoidal function as an activation function in all experiments if not specified.

### Network optimization

We formulate the compression task as an optimization problem

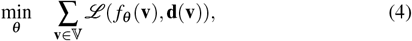

where 𝕍 is the grid coordinate system matching the input data volume; **d**(**v**) is the intensity value of original data at vertex **v**; *ℒ* measures the difference between decompressed data and the original version before compression. In the implementation, we adopted the L2 norm loss function in all experiments if not specified. During optimization, we use the Adaptive Moment Estimation Algorithm (Adam^62^) to update the network parameters in an iterative manner until convergence and apply mini-batch gradient descent to fit the limited memory by randomly sampling a batch of **v** from 𝕍. Users can flexibly adjust the compression quality by specifying training steps. Unless otherwise specified, the default value for training steps is set at 80,000.

### Adaptive partitioning

Since strong data correlation only exists locally, it is impractical to learn a unified implicit neural function to represent the whole data volume. Instead, here we decompose the whole data volume into blocks and learn their own implicit neural function separately. Although equal-sized partitioning is a commonly employed scheme for efficient data representation, it fails to extract reasonable within-block correlation in data with the intricate spatial distribution of information. To accommodate diverse biomedical data, we design an adaptive partitioning strategy, drawing inspiration from ACORN^35^. The strategy formulates data partitioning as an Integer Linear Programming (ILP) problem to find a proper tree structure organization of the original data volume. This strategy not only mitigates the substantial GPU memory overhead that ACORN incurs when dealing with large-sized data but also outperforms ACORN in terms of compression quality on smaller-sized data (Supplementary Figure 12).

For an *N*-dimensional visual data, let *S*_1_ × · · · × *S*_*N*_ denote its spatial coordinates and *c* denote the channels (e.g., color, spectrum, etc.), we can define an *H*-level tree data structure. At the first level, the root node can be described as 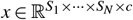. At finer levels, 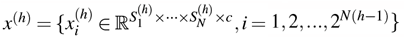 represents a set of blocks with 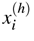 being the *i*th node at the *h*th level, and the dimension of the block is

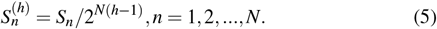

A partitioning scheme can be described as a tree with binary node values: 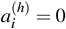 indicates the *i*th node at *h*th level being a leaf node with sufficiently high within-block correlation or 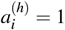 represents a non-leaf node needs further partitioning. Further, we define an objective as Eq. (6) to explicitly model the within-block correlation for pursuing a good partitioning scheme

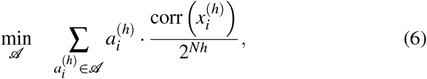

where corr(·), the within-block correlation (please refer to Supplementary Note 1 for details), is calculated by the proportion of 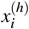s top Γ amplitude spectrum

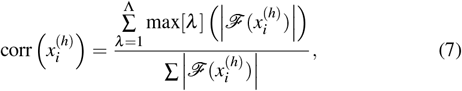

with function max[*λ*](·) returning the *λ* th largest entry and *𝒜* denotes all possible partitioning strategies

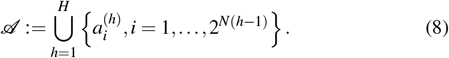

The feasible set of our partitioning strategy is given in Eqs. (9) and (10).

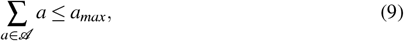

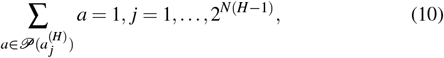

with *𝒫* (·) denoting a path to the root node. Here Eq. (9) defines the upper bound of the number of blocks which can be limited by a theoretical minimum number of parameters in a single implicit neural function to maintain representation capability^63^, and Eq. (10) is introduced according to the rules of the tree data structure. Based on the above formulation, searching for the best partitioning strategy is formulated into an ILP problem that can be solved efficiently by state-of-the-art solvers.

### Adaptive parameter allocation

Under the derived partitioning scheme, we further develop a heuristic parameter allocation strategy to allocate the given parameter budget across the blocks (Extended Data Fig. 3). Initially, the number of parameters allocated to each small block is proportional to 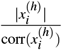, where 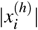 denotes the size of block 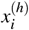 and 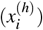 measures the within-block correlation. This fraction roughly reflects the complexity (or representation difficulty) of the block. Then, we alternatively optimize the implicit neural function of each block and adjust the parameter allocation according to the fitting loss on those blocks. On the one hand, we allocate more parameters to enhance the accuracy of the under-performing blocks with larger fitting loss. To be more specific, we assign the implicit neural function of under-performing blocks more neurons, and zero initialize the newly added weight and bias to secure consistency with the previous function,

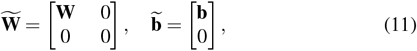

where **W** and **b** are respectively the weights and bias of the original model, and 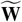 and 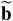 are the updated version. On the other hand, the out-performing blocks with low fitting loss tend to be pruned and quantized for higher sparsity (i.e., containing more zero values) by Condensa^64^. Such a strategy would facilitate higher decompression quality and more consistent fidelity over the whole data.

### Regulating the compression ratio

For our network architecture (Extended Data Fig. 1), the number of neurons in the input layer *M*_*in*_ is determined by the spatial dimensions of the original data, and that in the output layer *M*_*out*_ by the channels, so the total number of parameters in the neural network *P* is determined by the number of hidden layers *L* and the number of neurons contained in each hidden layer *m* as

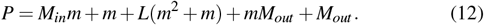

Then the size of the compressed data *P*_*s*_ can be calculated as *P*_*s*_ = *P* × *bits*, where *bits* is related to the numerical precision of neurons, e.g., *bits* = 32 for single-precision and *bits* = 16 for half-precision.

To regulate the compression ratio *r*, BRIEF first calculates the file size of compressed data (i.e., the neural network representation) as *P*_*s*_ = *F*_*s*_*/r*, where *F*_*s*_ is the size of original data. Correspondingly, the number of parameters can be derived as *P* = *P*_*s*_*/bits*. Given an *L*-layer network structure, BRIEF can rewrite Eq. 12 into a univariate quadratic equation

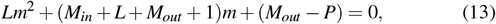

whose closed-form integral solution is

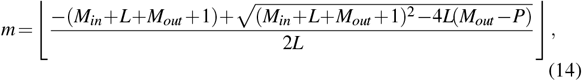

where ⌊·⌋ means rounding down operation. Finally, BRIEF, following this workflow, will approximately achieve the given compression ratio *r*, with only slight deviation coming from the facts that the number of neurons can only be integral and some extra bytes are required for the side information. A detailed schematic diagram is shown in Extended Data Fig. 4.

### Prioritizing fidelity of user defined ROIs

Our approach supports prioritizing compression quality for different regions. This advantageous feature is attributed to the fact that we formulate the compression task as an optimization problem defined in Eq. (4), which can easily incorporate region specific importance to guide the compression process. Specifically, one can generate a weight *ω*(**v**) over the grid spatial coordinate 𝕍, with a greater value indicating higher fidelity in compression. Then the original optimization problem can be modified into

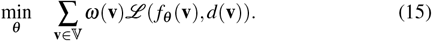

Such user-friendly compression is quite useful but not limited to the following cases: (i) Avoiding global performance degeneration caused by a small number of extreme values (e.g., with magnitudes higher or lower intensity) by excluding data fidelity at these positions. Otherwise, ensuring the fidelity of these extreme portions might harm the overall compression quality largely; (ii) Making extensive use of the budget, by giving high priority to the ROIs, which is especially important when facing a severe bandwidth or storage shortage.

### Quantitative fidelity criteria

For a 2D image of size *N*_*W*_ ×*N*_*H*_, suppose Φ(*x, y*) and Ψ(*x, y*) represent the original data and the decompressed version respectively, the fidelity of the decompressed data are evaluated in terms of widely used criteria, including peak signal-to-noise ratio

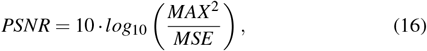

with *MAX* being the dynamic range of the original data, *MSE* being the mean squared error

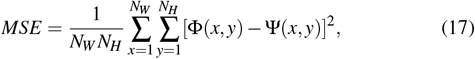

and structural similarity index

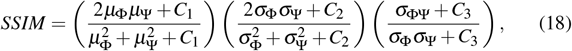

with *μ*_Φ_ and *μ*_Ψ_ denoting the mean of Φ and Ψ respectively, *σ*_Φ_^2^ and *σ*_Ψ_^2^ denoting the variance of Φ and Ψ respectively and *σ*_ΦΨ_ denoting the covariance between Φ and Ψ. For the data the morphological structure other than actual intensity values matters, such as the neurons and vessels, we first binarize Φ and Ψ using a threshold *τ* to obtain Φ_*τ*_ and Ψ_*τ*_, where 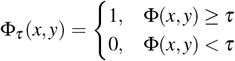 and the same for Ψ_*τ*_ (*x, y*). *τ* represents the threshold determining whether a pixel region corresponds to a neuron/vessel or belongs to the background. Then we adopt the indicator F1-score from the front-background segmentation task to evaluate the decompressed data structure

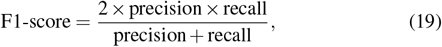

with precision = *TP/*(*TP* + *FP*), and recall = *TP/*(*TP* + *FN*). Since this indicator is related to the threshold *τ* used in the binarization process, we use F1(*τ*) to abbreviate this indicator.

For 3D volume, the fidelity score is calculated by taking an average over all the slices (2D images).

### Collection of biomedical data

Neurons and vessel data were imaged by our self-developed HD-fMOST system^9^. Please refer to Supplementary Note for details on sample preparation and HD-fMOST system. Organic CT data were downloaded from HiP-CT project^2^. XNH, MRI, Fundus, Calcium imaging, bright-field microscopy, two-photon microscopy, and confocal microscopy data were released from the published articles^8,53–57,65,66^.

### Downstream vision tasks

To provide biology-relevant evidence that BRIEF, in comparison to state-of-the-art algorithms, can retain more signals of interest, we perform downstream tasks on both the original and decompressed datasets. The results show that one can achieve comparable performance before and after applying BRIEF compression, but suffers from accuracy degeneration when using other compressors. Therefore, we can conclude that BRIEF can serve as a reliable compressor for biomedical data.

For neurons, we used an open source software Gtree^3^ to locate the nerve fibers with 3D tracing. We adjusted the threshold parameter for each data to make the tracking results more continuous. For quantitative evaluation, the F1-score is used to numerically measure the differences in tracing results before and after compression. To avoid ambiguity, we use the Reconstruction F1-score to denote this metric. For the tracing points of decompressed data, the ones having a counterpart within 4*μm* in the tracing result of original data are counted as *TP*, otherwise counted as *FP*. Conversely, the tracing points in original data without correspondence within its 4*μm* neighborhood in the decompressed counterpart are regarded as *FN*. Since the Reconstruction F1-score assesses the compression quality via measuring the consistency of the 3D tracing results on the decompressed data and original version, the Reconstruction F1-score of the original data equals 1 and a closer-to-1 score implies higher data fidelity.

For vessels, we used Imaris 9.0 software to reconstruct their 3D vascular structures with intensity ranging from 20 to 2000. For rendering, we choose the Blend mode which uses all the intensities along the viewing direction together with their transparency to produce the final 3D visual demonstration. For organic CT, the ROIs were segmented using marching squares^50^ and intelligent scissors^51^ semi-automatically by experts with anatomical knowledge and expertise in diagnosis.

For the retrieval of calcium activity traces of individual neurons, we simply took the average over the intensities within a window around each neuron.

### Workstation configuration

All of the mentioned experiments are conducted on a workstation with AMD Ryzen Threadripper 3970X Processor and NVIDIA GeForce RTX3090 GPUs. We use PyTorch and CUDA to implement our compression algorithm, and Gurobi to solve related ILP problems.

### Serialization of neural networks

For a given data, we need to save its partitioning information and organize the compressed file by creating a folder for each block, named by its position and size, e.g., for a 16× 128× 128 block starting from(0, 256, 0) is named “0 15-256 383-0 12”. After compressing a block (i.e., optimizing its implicit neural function), we serialize both its parameters and side information and save them into a folder. As for the network parameters, we only store the weight and bias of each layer as a binary file named with its layer index. The side information, including descriptions of the network structure, bit depth, and size of the original data, and inverse normalization coefficients, are saved as a YAML file. This file can be very small since all the items are discrete data, i.e., being countable and can only take certain values, and we can simply build a table listing all possible values and record the index. In addition, the length of the above file name is much smaller than the limit of modern file systems.

### Data preprocessing

Before optimization, we normalize the coordinate vector **v** within range [− 1, 1] and the intensity **d**(**v**) within range [0, 100] by Min-Max normalization. The original range and the normalization method are recorded in the side information which will be used in decompression.

### Data postprocessing

During decompression, we need to combine the separately decompressed blocks into whole data. As a block-wise compressor, there might exist artifacts along the partitioning boundary, especially at high compression ratios. To address this issue, we adopt an adaptive deblocking filter^67^ to adaptively identify the boundaries that need to be smoothed and perform smoothing based on the values near the boundaries (please refer to Supplementary Note 2 for details).

## Data availability

Neurons and vessel data are available from the corresponding author on reasonable request. Organic CT data are available from HiP-CT project^4^. XNH, MRI, Fundus, Calcium imaging, bright-field microscopy, two-photon microscopy, and confocal microscopy data are available from the published articles^8,53–57,65,66^.

## Code availability

The CUDA implementation of BRIEF is publicly available at https://github.com/RichealYoung/BRIEF_CUDA, and the PyTorch version is released at https://github.com/RichealYoung/BRIEF_PyTorch.

## Acknowledgements

We acknowledge Prof. T. Xia for building collaborations among several groups. We thank Dr. F. Bao and Dr. Q. Cao for their valuable suggestions on the manuscript. This work is jointly supported by the Ministry of Science and Technology of the People’s Republic of China (Grant No. 2020AAA0108202), Beijing Municipal Natural Science Foundation (Grant No. Z200021), and the National Natural Science Foundation of China (Grant Nos. 61931012, 62088102).

## Author contributions

J. S., R. Y., T. X., and Y. C conceived this project. Q. D., Q. L., and J. S. supervised this research. R. Y. and T. X. designed the BRIEF architecture. T. X., R. Y., and Y. C. designed the adaptive partitioning and allocation algorithms. Y. C., T. X., A. L., J. Q., and J. W. conducted the experiments and data analysis. A. L., X. W., R. L., S. B., Y. C., and T. X. collected the biomedical data. All the authors participated in the writing of this paper.

## Competing interests

The authors declare no competing financial interests.

**Extended Data Fig. 1.**
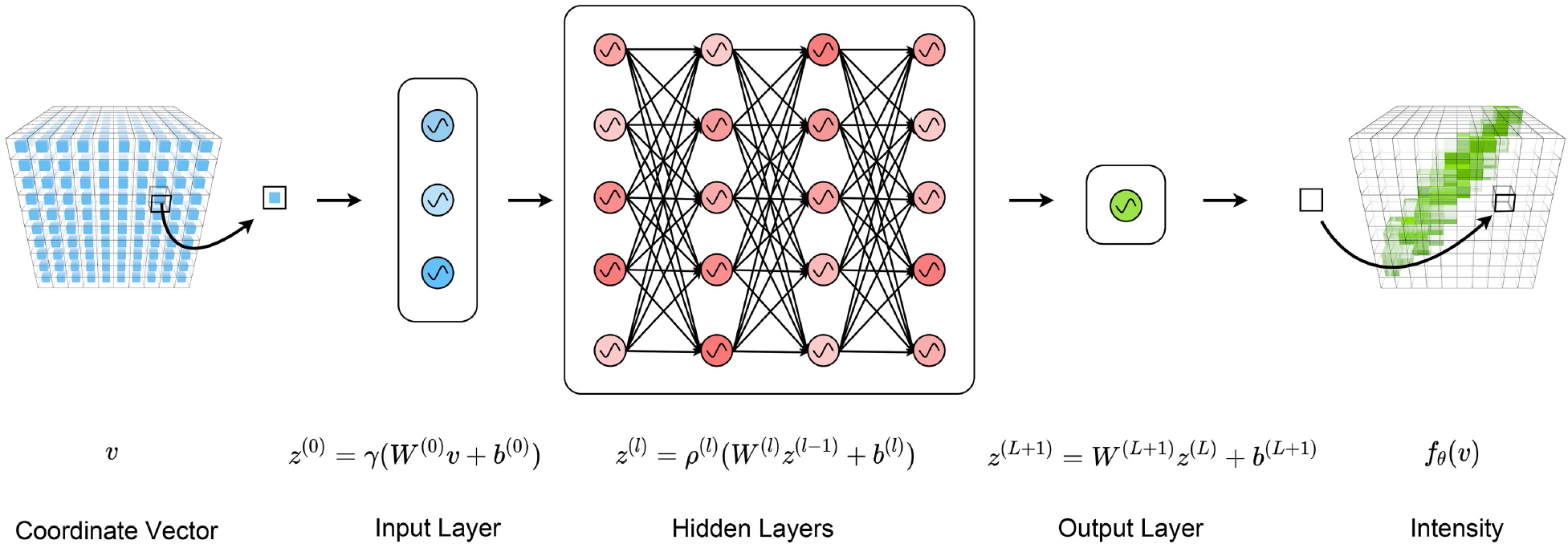
A visualization of our neural network architecture. We use a SIREN-like MLP as our neural network. The input *v* is the coordinate vector of each voxel in the original data. Then *v* is embedded into a higher dimensional space using a sinusoidal function *γ* at the input layer. The hidden layers, a fully connected neural network with activation function *ρ*, model the underlining data structure *z*^(*L*)^with the embedding *z*^(0)^. The output layer finally maps *z*^(*L*)^ into the corresponding voxel’s intensity, *f*_*θ*_ (*v*), written as the function of coordinate vector *v*. The numbers of neurons in the input layer and output layer signify the coordinate’s dimension and the number of channels of original data respectively. The number of neurons in hidden layers can be adjusted freely according to the need.

**Extended Data Fig. 2.**
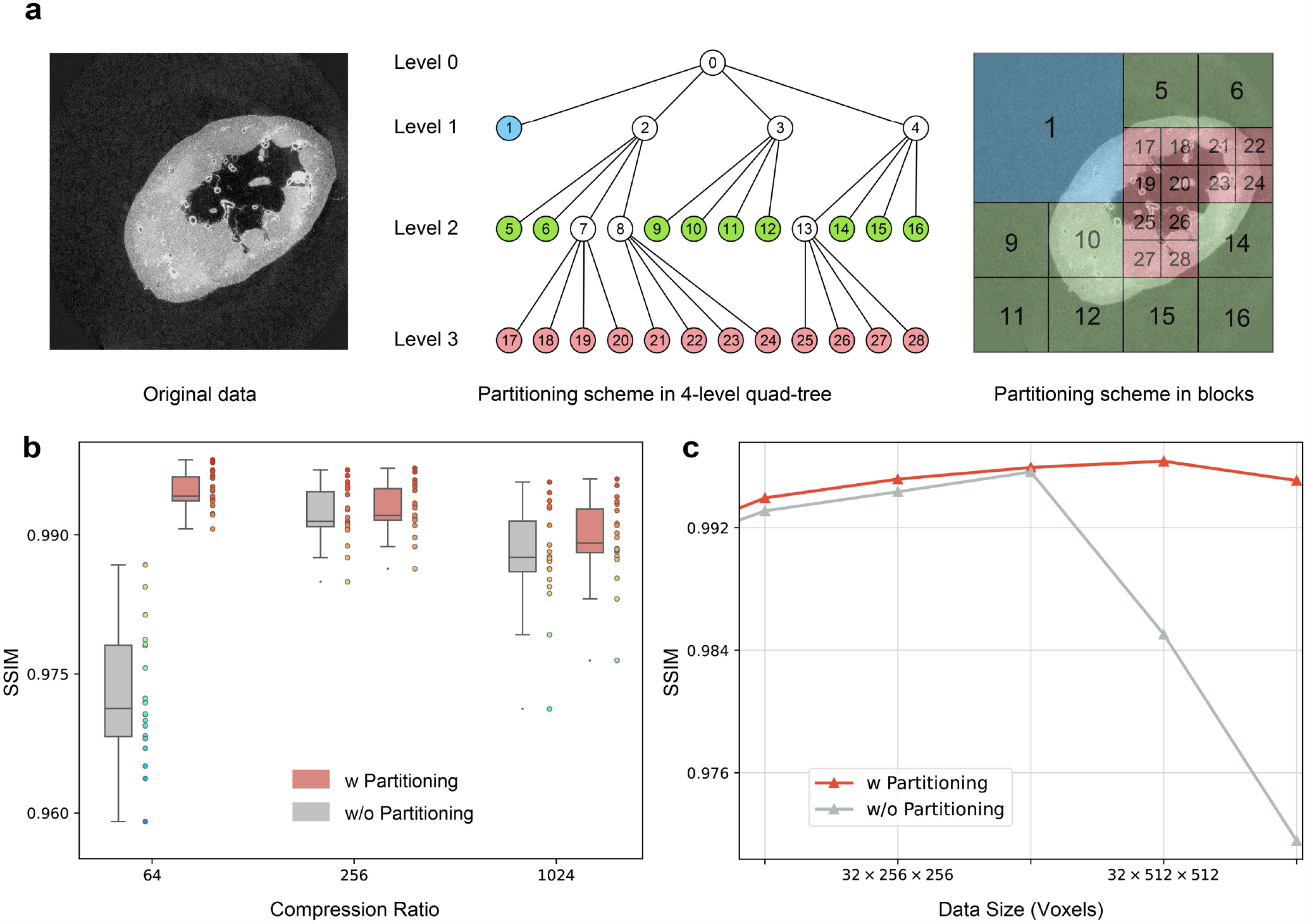
The adaptive partitioning. **a**, Schematic diagram of adaptive partitioning for two-dimensional data (left) based on 4-level quad-tree. The optimal solution of Eq. (6) by ILP solver is shown both in tree-view (middle) and blocks-view(right). The leaf nodes with sufficiently high within-block correlation are filled with different colors according to their levels, while the non-leaf nodes line to their further partitioned nodes. The numbers and colors on the blocks (right) correspond to the ones on the nodes (middle). **b**,**c**, The ablation study of adaptive partitioning at different compression ratios and data sizes. We conduct experiments under three compression ratios (64 ×, 256 ×, 1024 ×) and five data size (16 ×128 ×128, 32 ×256 ×256, 32 × 256 ×512, 32 ×512 ×512, 64 × 512 × 512) on tens of organic CT data. The compression quality is measured by SSIM. Results show that the adaptive partitioning strategy improves performance especially when the compression ratio is relatively low and the data size is relatively large since the neural network representing the whole data (i.e., without partitioning) has too many parameters to be optimized.

**Extended Data Fig. 3.**
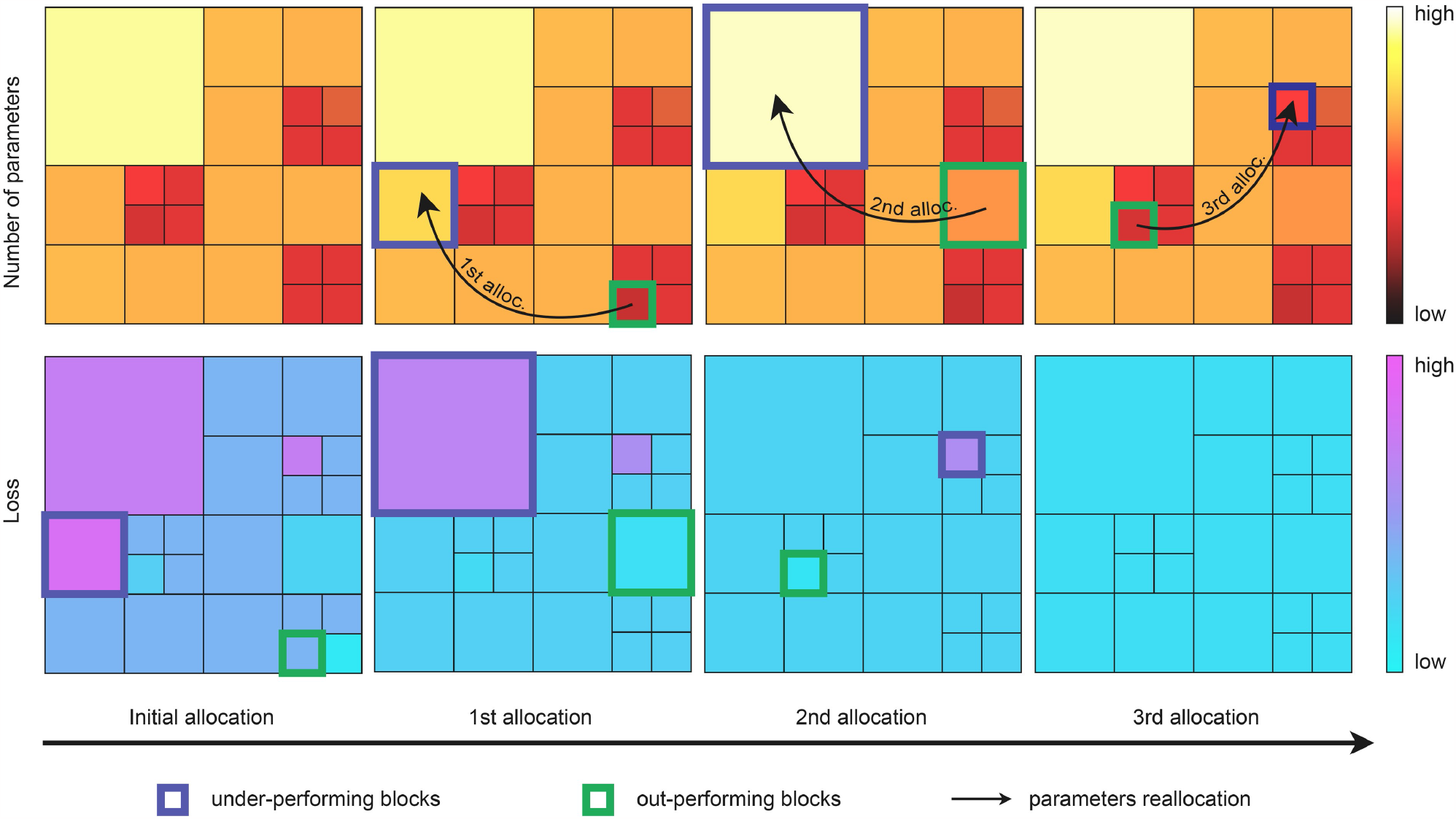
The adaptive parameter allocation. BRIEF initially allocates the number of parameters according to each block’s size and within-block correlation (top left). After certain steps of fitting, the fitting loss of each block appears to vary due to different fitting difficulties (bottom left). Then BRIEF reallocates the number of parameters from the out-performing blocks (highlighted in the blue box) to under-performing blocks (highlighted in the green box) and fits those blocks again. BRIEF repeats reallocation and fitting until certain times or the loss are evenly distributed. The numbers of parameters and fitting loss are color-coded.

**Extended Data Fig. 4.**
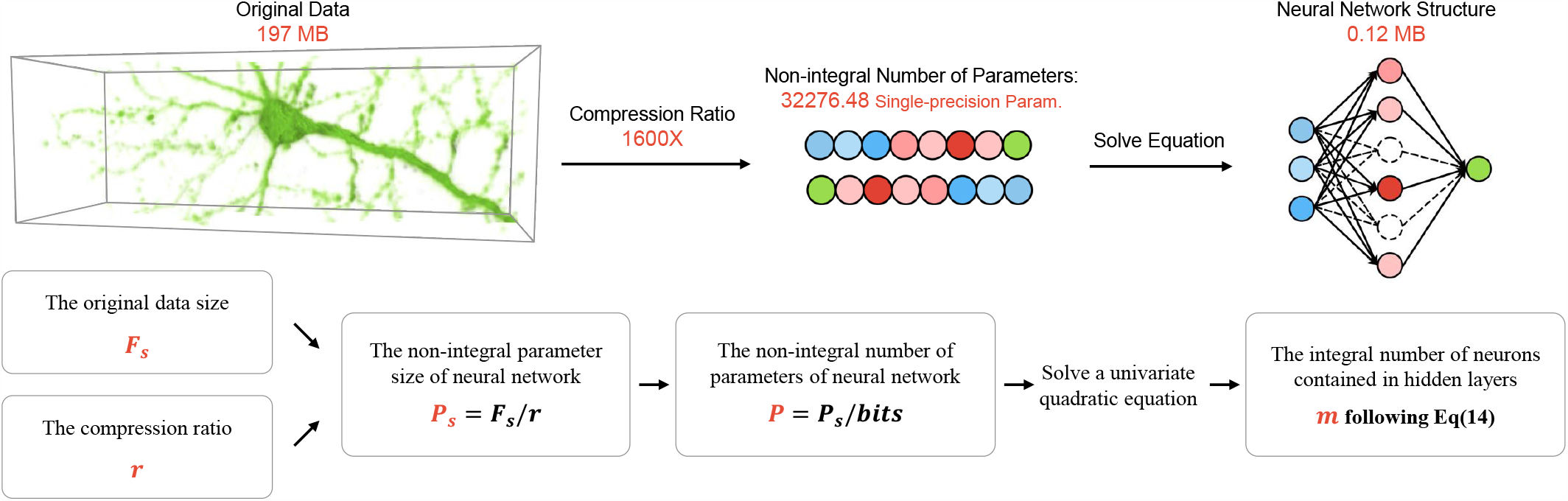
Flow chart of regulating compression ratio. For a compression task, given the original data size *F*_*s*_ and compression ratio *r*, the non-integral parameter size of a neural network can be calculated as *P*_*s*_ = *F*_*s*_*/r*. And the non-integral number of parameters will be *P* = *P*_*s*_*/bits*. The *bits* of a single-precision parameter is 32 and of a half-precision one is 16. By solving the univariate quadratic equation in Eq. (13), BRIEF can derive the feasible integral number of neurons contained in hidden layers through the closed-form solution given in Eq. (14).

## Footnotes

1 https://github.com/GTreeSoftware/GTree

2 https://human-organ-atlas.esrf.eu/

3 https://github.com/GTreeSoftware/GTree

4 https://human-organ-atlas.esrf.eu/

