## Supplementary Materials for "Sharing Massive Biomedical Data at Magnitudes Lower Bandwidth Using Implicit Neural Function"

#### Contents

**1 Within-block Correlation in Adaptive Partitioning 2**
**2 Adaptive Deblocking Filter 2**
**3 Additional Benefits of Noise Suppression 3**
**4 Sample Preparation and Imaging Setup of 3D Mouse Brain Data 3**
**5 Supplementary Figures 4**
**6 Supplementary Tables 13**
**References 14**

### 1 Within-block Correlation in Adaptive Partitioning

Strong correlation exist only in local regions of the entire biomedical data, so the divide-and-conquer strategy (i.e., partition data into blocks and conduct block-wise encoding) is an advantageous option for efficient data compression. Although simple uniform partitioning is widely used in commercial methods like H.264[1] and H.265[2], a deep understanding of mechanisms behind the advantages of partitioning for data representation can shed more light on designing better partitioning scheme. To address this issue, we offer a novel perspective that theoretically analyzes how partitioning advances BRIEF.

Here we explain on a three-layer BRIEF architecture without bias $\mathbf{b}$ for convenience and it can be extended to deeper networks. The first layer in BRIEF conducts frequency mapping for the input coordinate vector $\mathbf{v}$, and the output $\mathbf{z}^{(0)}$ can be described as

$$\mathbf{z}^{(0)} = \sin\left(\mathbf{W}^{(0)}\mathbf{v}\right) = \sin(\Omega\mathbf{v}), \tag{1}$$

in which $\mathbf{W}^{(0)}$ is the weighting matrix, and we replace $\mathbf{W}^{(0)}\mathbf{v}$ with $\Omega\mathbf{v}$ to represent the input frequencies. The second layer takes the linear combination of above sine wave components at varying mapping frequencies as input, with $\mathbf{W}^{(1)}$ denoting the combination coefficients of higher order terms of the input frequencies, followed by a sinusoidal activation function. The output of the $n$th neuron in the second layer can be written as:

$$\mathbf{z}_n^{(1)} = \sin(\mathbf{W}_n^{(1)}\sin(\Omega\mathbf{v})). \tag{2}$$

Using the Euler formula we have Eq. (3), which further turns into Eq. (4) after Fourier series expansion, in which we introduce the Bessel function to exhibit the weights of input frequencies and their high-order harmonics. By further derivation, one can get Eq. (4) in which $t_k\Omega_k$ represents a linear combination of integer harmonics of the input frequencies, indicating expression capability of the BRIEF architecture for a wide range of frequencies.

$$\mathbf{z}_n^{(1)} = \mathrm{Im}\left\{\prod_{k=0}^{K-1} exp\left(j\mathbf{W}_{n,k}^{(1)}\sin(\Omega_k\mathbf{v})\right)\right\} \tag{3}$$

$$= \mathrm{Im}\left\{\sum_{t_0=-\infty}^{\infty}\cdots\sum_{t_{K-1}=-\infty}^{\infty}\prod_{k=0}^{K-1} J_{t_k}(\mathbf{W}_{n,k}^{(1)})\exp(jt_k\Omega_k\mathbf{v})\right\} \tag{4}$$

$$= \sum_{t_1,\ldots,t_K=-\infty}^{\infty}\left(\prod_{k=0}^{K-1} J_{t_k}(\mathbf{W}_{n,k}^{(1)})\right)\sin\left(\sum_{k=0}^{K-1} t_k\Omega_k\mathbf{v}\right). \tag{5}$$

Considering the decaying behavior of Bessel functions $J_{t_k}(\mathbf{W}_{n,k}^{(1)})$, the high order harmonics in $\mathbf{z}_n^{(1)}$ tend to have smaller weights than the input frequencies. Further, the third layer is again a linear combination of the output $\mathbf{z}_n^{(1)}$ given in Eq. (6), and so forth for a deeper network:

$$f_\theta(\mathbf{v}) = \mathbf{w}^{(2)\top}\mathbf{z}^{(1)} = \mathbf{w}^{(2)\top}\sin(\mathbf{W}^{(1)}\sin(\Omega\mathbf{v})), \tag{6}$$

with $\top$ denotes matrix transpose.

On the whole, the above BRIEF architecture acts as an inductive bias mechanism, which focuses most energy of the output signal in a narrow band around the input frequencies $\Omega$. Considering such frequency concentration property, BRIEF is more suitable for representing local signals which are of much more concentrated frequency features. In other words, a very small number of input frequencies would cover the spatial spectrum of a local area and of high representation accuracy, while far insufficient for large data and thus cause severe reconstruction artifacts.

Inspired by the above analysis, we utilize the frequency concentration property of BRIEF and propose to use the ratio of data $x$'s top $\Gamma$ amplitude spectrum

$$\mathrm{corr}(x) = \frac{\sum_{\gamma=1}^{\Gamma}\max[\gamma](|\mathscr{F}(x)|)}{\sum|\mathscr{F}(x)|}, \tag{7}$$

to represent the within-block correlation, with the function $\max[\gamma](\cdot)$ denoting the $\gamma$th largest item in the input vector. This score theoretically infers a good partitioning for efficient block-wise BRIEF representation, under a given compression ratio.

### 2 Adaptive Deblocking Filter

During decompression, we need to combine the separately decompressed blocks into whole data. As a block-wise compressor, there might exist artifacts along the partitioning boundary, especially at a high compression ratio. To address this issue, we adopt an adaptive deblocking filter[3] to adaptively identify the boundaries that need to be smoothed and perform smoothing based on the values near the boundaries. The deblock filtering step was not included in the decompression time in this paper, and it takes less than 0.1 seconds on our workstation. Denote the intensities along a line across two neighboring blocks as $p_3$, $p_2$, $p_1$, $p_0$, $q_0$, $q_1$, $q_2$, $q_3$, with the boundary lying between $p_0$ and $q_0$, as illustrated in Supplementary Figure 13. To avoid excessive smoothing and loss of details , we first identify the boundaries that need to be smoothed as those meeting the following three conditions simultaneously

$$|p_0 - q_0| < \alpha(Index_A), \tag{8}$$

$$|p_1 - p_0| < \beta(Index_B), \tag{9}$$

$$|q_1 - q_0| < \beta(Index_B), \tag{10}$$

where $\alpha$ and $\beta$ are defined as

$$\alpha(x) = 0.8(2^{x/6} - 1), \tag{11}$$

$$\beta(x) = 0.5x - 7. \tag{12}$$

Here the value ranges of $Index_A$ and $Index_B$ are both $[0, 51]$ empirically. Intuitively, a small $Index_A$ or $Index_B$ helps to preserve more details, while a larger setting would remove blocking artifacts better but at risk of losing some details. In our implementation, we set filter parameters adaptively according to the partitioning strategy which reflects the information distribution of the original data.

If all the above three conditions are met, the intensities along the line will be updated as

$$p_0^{'} = p_0 + \Delta_0, \tag{13}$$

$$q_0^{'} = q_0 - \Delta_0, \tag{14}$$

$$p_1^{'} = p_1 + \Delta_{p1}, \tag{15}$$

$$q_1^{'} = q_1 + \Delta_{q1}, \tag{16}$$

where $\Delta_0$, $\Delta_{p1}$ and $\Delta_{q1}$ are calculated as

$$\Delta_0 = (4(q_0 - p_0) + (p_1 - q_1) + 4)/8, \tag{17}$$

$$\Delta_{p1} = (p_2 + (p_0 + q_0 + 1)/2 - 2p_1)/2, \tag{18}$$

$$\Delta_{q1} = (q_2 + (q_0 + p_0 + 1)/2 - 2q_1)/2. \tag{19}$$

To avoid blurring, we constrain the value range of $\Delta_0$ within $[-c_0,c_0]$, and $\Delta_{p1}$, $\Delta_{q1}$ within $[-c_1,c_1]$

$$\Delta_0 = \min(\max(-c_0, \Delta_0), c_0), \tag{20}$$

$$\Delta_{p1} = \min(\max(-c_1, \Delta_{p1}), c_1), \tag{21}$$

$$\Delta_{q1} = \min(\max(-c_1, \Delta_{q1}), c_1), \tag{22}$$

where $c_0$ and $c_1$ are used to limit the smoothing effect. We set $c_0 = 20, c_1 = 20$, while increasing $c_0$ by 1 if the following conditions are met.

$$|p_2 - p_0| < \beta(Index_B), \tag{23}$$

$$|q_2 - q_0| < \beta(Index_B). \tag{24}$$

#### 3 Additional Benefits of Noise Suppression

The recorded biomedical data might suffer from system noise, typically shot noise, which would cause artifacts after lossy compression. Because BRIEF models the intrinsic structure of the target data, it would naturally suppress the noise that are statistically of different spatial distributions from the true signals. To demonstrate this additional benefit, we conduct compression using BRIEF and H.265 on tens of 3D vascular data with different simulated shot noise, which follows a Poisson distribution with parameter $\lambda$ varying from 0 to 1000. The comparison is shown in Supplementary Figure 4. From the results, one can see that H.265's compression quality degrades dramatically with the increase of noise level, while BRIEF maintains a high compression quality. In addition, after being contaminated by noise, the vessel in H.265's decompressed version is incorrectly broken, while BRIEF exhibits superior noise robustness.

#### 4 Sample Preparation and Imaging Setup of 3D Mouse Brain Data

The Neural and Vascular data volumes in this article are both captured by fluorescent imaging of a fixed mouse brain. An 8-week-old Tek0-Cre*Ai47 mouse and a 4-month-old C57BL/6J mouse were used in this study. The mice were housed on a 12-hour light/dark cycle with food and water ad libitum. The Tek0-Cre*Ai47 mouse was generated by crossing Tek0-Cre transgenic mice and GFP Cre reporter Ai47 mice for obtaining whole-brain vascular labeling. To identify projections from the motor cortex to the fastigial nucleus in the C57BL/6J mouse, AAVretro-CAG-Cre was injected into the latter region and AAV9-CAG-Flex-GFP was injected into the former, respectively. PI staining was used for obtaining cytoarchitectonic information, which was essential for determining the locations of brain structures. Brains were embedded in HM20 for whole-brain imaging. All experimental procedures were approved by the Animal Experimentation Ethics Committee of Huazhong University of Science and Technology.

We used the high-definition fluorescent micro-optical sectioning tomography (HD-fMOST) system to perform whole-brain dual-color imaging at a voxel size of $0.32\times0.32\times1\mu m^3$. The raw file size of the whole-mouse-brain data of the Tek0Cre*Ai47 mouse is ∼2.82 TB, containing ∼12,000 coronal slices with 29005×24486 pixels for each. The whole-brain data of the C57BL/6J mouse occupies ∼10.9 TB, containing 5,160 coronal slices, each with 28692×20005 pixels.

#### 5 Supplementary Figures

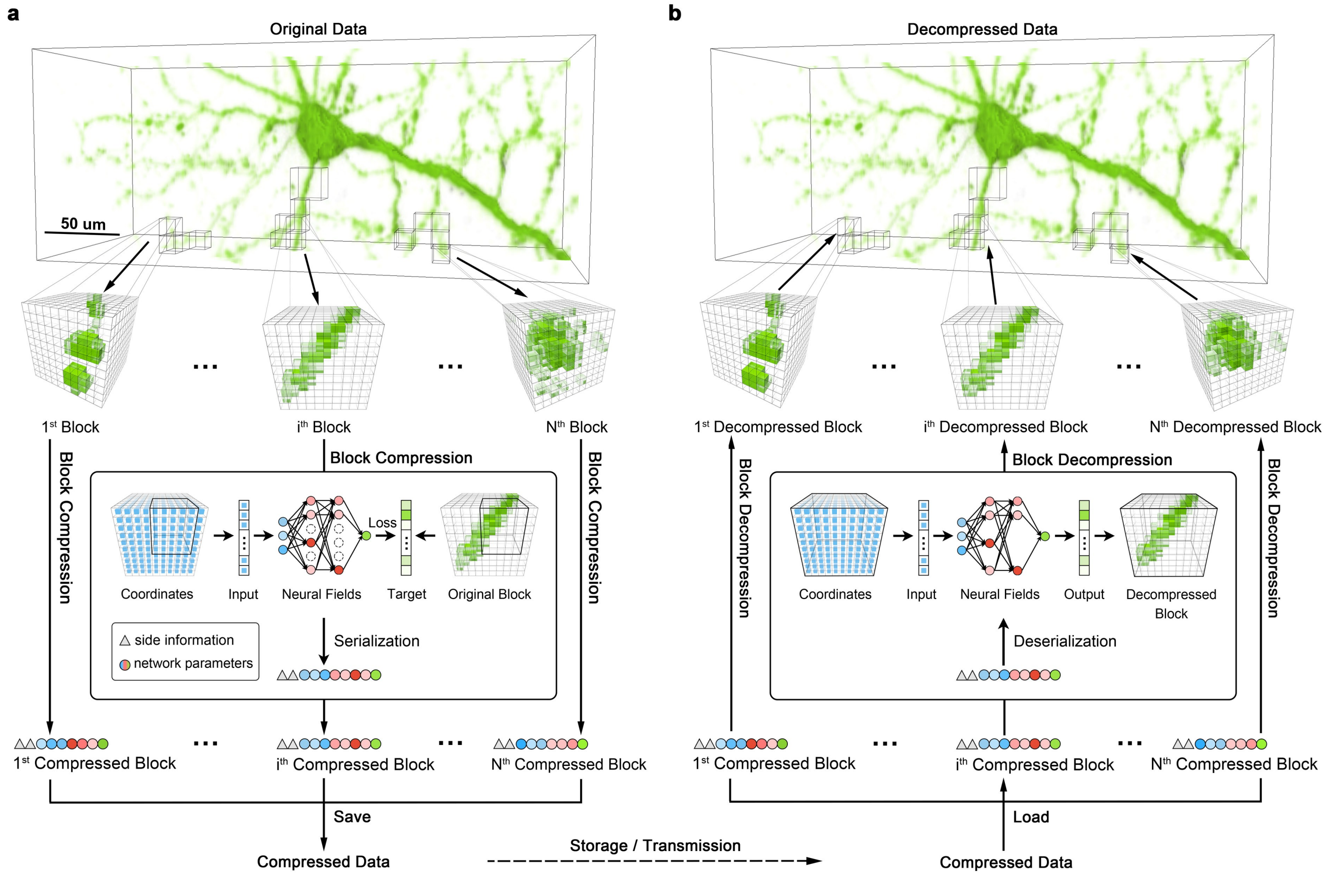

**Supplementary Figure 1. Diagram of the compression and decompression stages of BRIEF. a**, During compression, the original data is firstly partitioned into blocks with similar complexity adaptively. Then each block undergoes separate compression via pursuing a neural network that maps a voxel coordinate to its corresponding intensity value and is encoded into a sequence of bits via serializing this best-matching network. During block-wise compression, each neural network fits the corresponding data block by optimizing and pruning its parameters. Finally, the serialized codes of all the blocks are concatenated together into a long-bit sequence and saved in a file called compressed data, which can be stored efficiently or transmitted at low bandwidth. **b**, To decompress the volumetric data, the holistic serialized sequences are firstly parsed into codes for different blocks, i.e., some neural networks, which can reconstruct corresponding data blocks via simple forward propagation. All the reconstructed blocks are then stitched together to form the whole data volume.

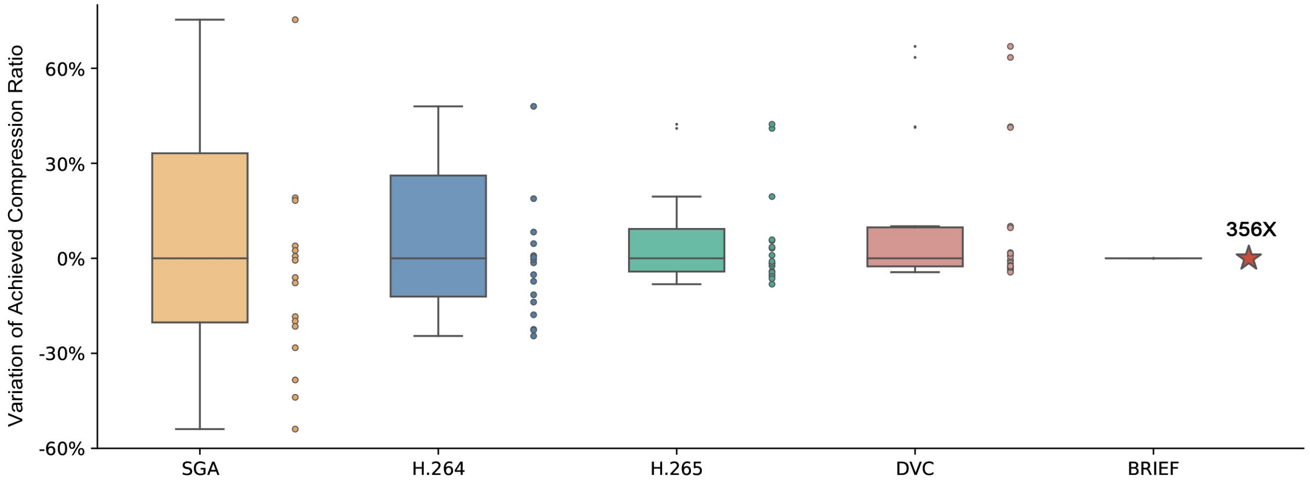

**Supplementary Figure 2. BRIEF supports precise specification of compression ratio.** Box plots of the final achieved compression ratio for a specified fixed target compression ratio around 356×, calculated from results on 16 randomly selected data volumes. Variation of achieved compression ratio indicates the degree of change in each achieved compression ratio compared to the median value. Only BRIEF can exactly achieve the target compression ratio (red star) for all the test data.

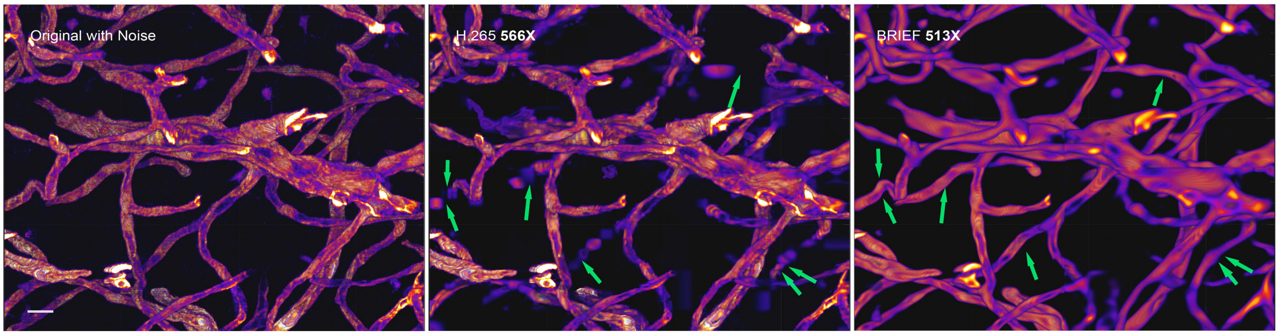

**Supplementary Figure 3. The comparison between BRIEF and H.265 on reconstruction of 3D vascular structure.** The 3D reconstruction of vascular structure in the original noisy vessel data (left), after applying compression by H.265 (middle) and BRIEF (right) at around 512× compression ratio. The capture of vessel data suffers from noise, resulting in a gray background and rough surface in the original data. Green arrows highlight the positions H.265 suffers from missing details while BRIEF achieved high fidelity reconstruction. Scale bar: $10\mu m$.

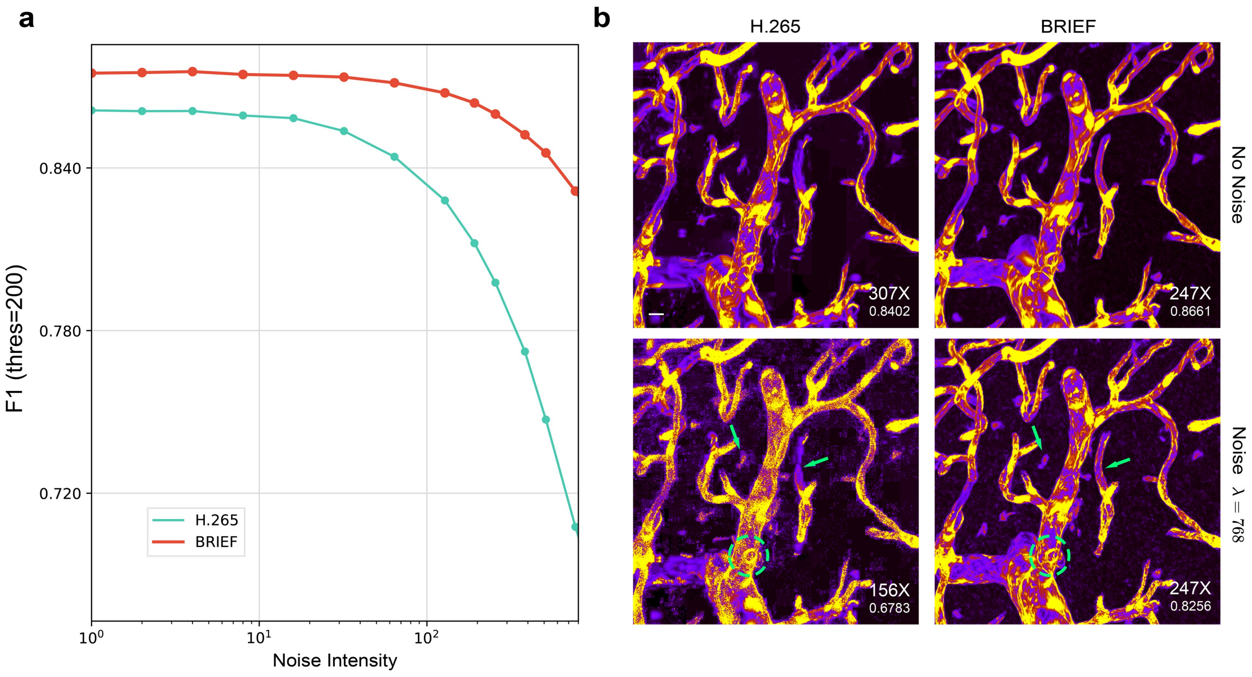

**Supplementary Figure 4. Noise suppression effect. a**, The comparison of the accuracy of a reconstructed 3D vascular structure after compression by H.265 and BRIEF, with F1 threshold being 200 and at ten different noise levels. The noise follows a Poisson distribution with parameter $\lambda$. Compared to H.265, whose compression quality degrades dramatically with the increase of noise level, while BRIEF exhibits superior noise suppression effect. **b**, The maximum intensity projection of the vessel data after applying compression by H.265 (left) and BRIEF (right) with (bottom) and without noise (top). The compression ratio and F1 with threshold 200 are labeled at the right bottom. The green arrows and circle highlight the regions where H.265 is disturbed by noise. Scale bar: $9\mu m$.

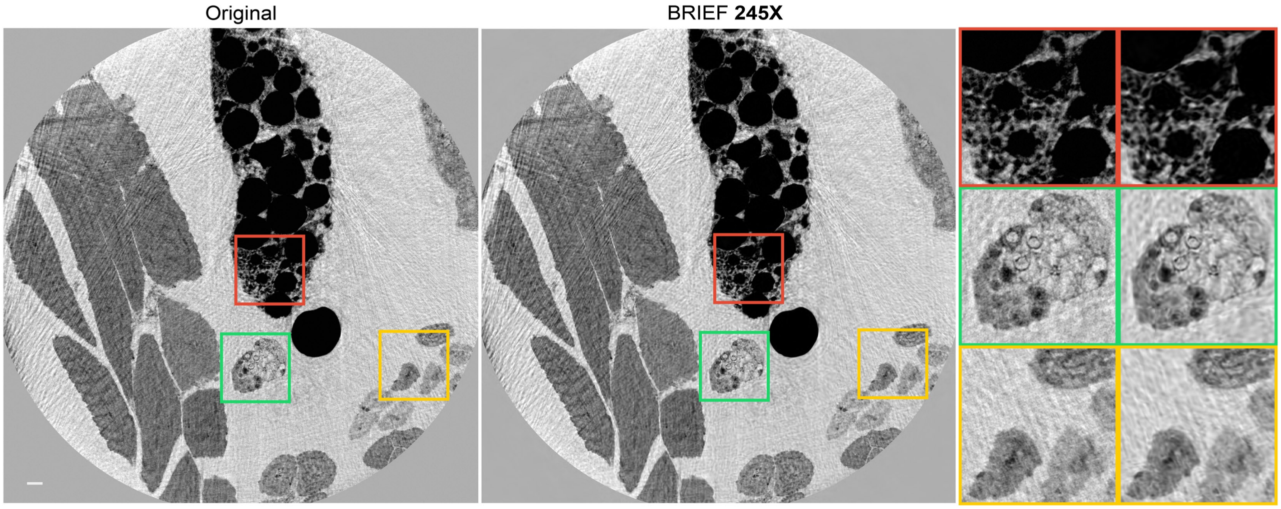

**Supplementary Figure 5. BRIEF's application on 3D volumetric data of drosophila leg captured by X-ray holographic nano-tomography (XNH).** The comparison between the original and BRIEF's decompressed data for XNH data volumes at around 245× compression ratio, with a zoomed-in view on the right side for clearer demonstration. The 3D volumes were released from the published article[4]. Scale bar: 20$\mu m$.

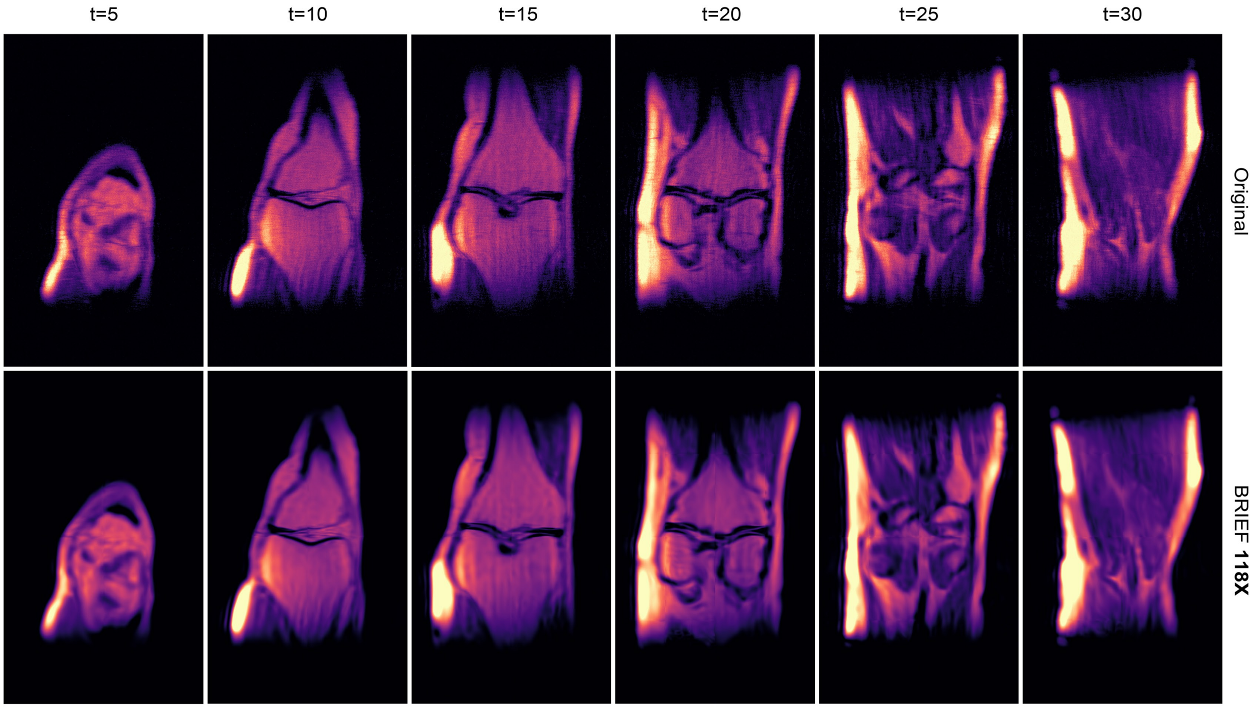

**Supplementary Figure 6. BRIEF's application on knee MRI data.** The comparison between the original and BRIEF's decompressed data for knee MRI data volumes at around 118× compression ratio. The 3D volumes were released from the published article[5].

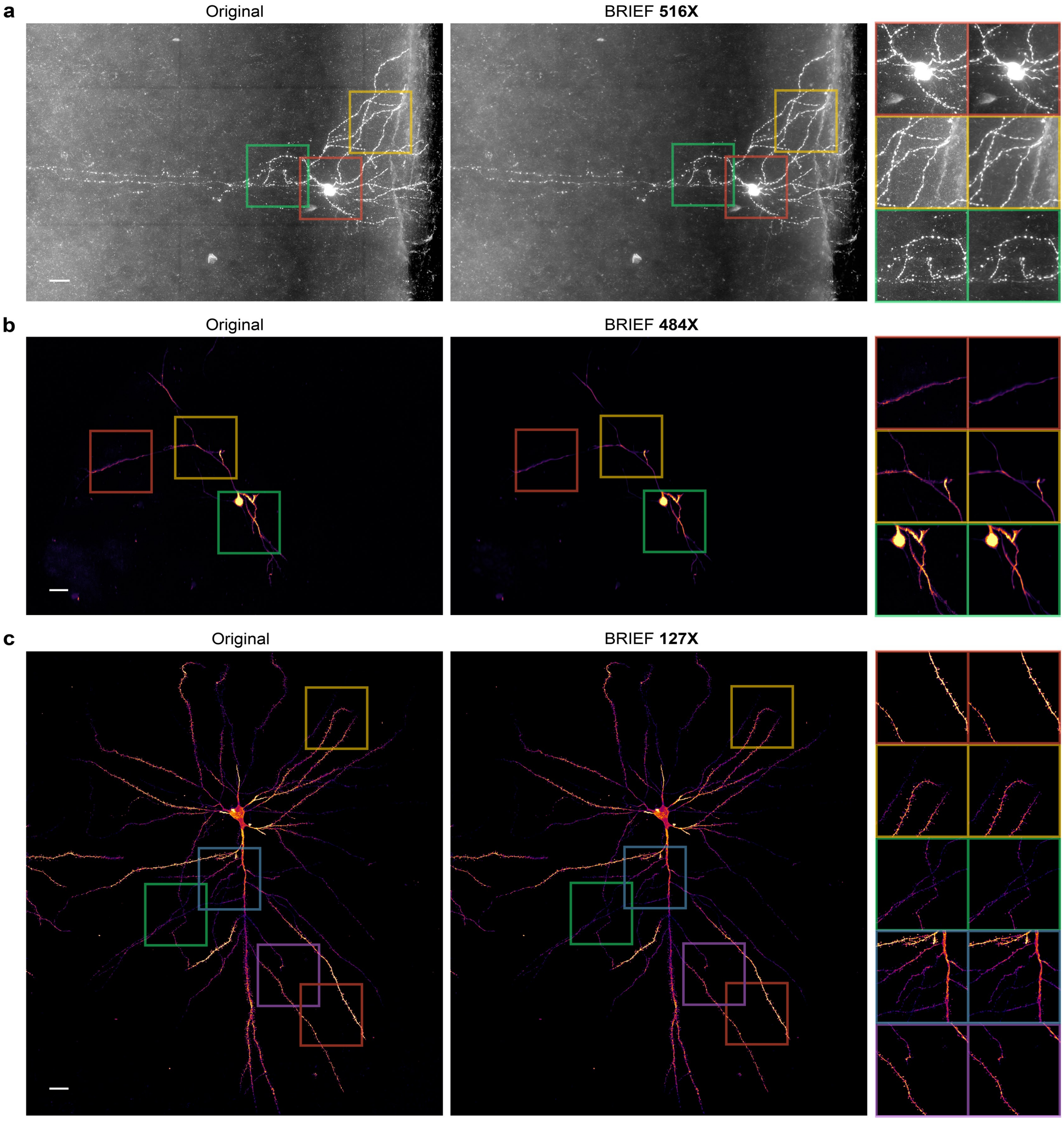

**Supplementary Figure 7. BRIEF's applications on brain neuronal data captured by bright-field, two-photon, and confocal microscopy.** **a**, The comparison between the original and BRIEF's decompressed data for brain neuronal data captured by bright-field microscopy[6] at around 516× compression ratio, with zoomed-in view on the right side for clearer demonstration. Scale bar: 20$\mu m$. **b**, The comparison between the original and BRIEF's decompressed data for brain neuronal data captured by two-photon microscopy[7] at around 484× compression ratio, with zoomed-in view on the right side for clearer demonstration. Scale bar: 20$\mu m$. **c**, The comparison between the original and BRIEF's decompressed data for brain neuronal data captured by confocal microscopy[8] at around 127× compression ratio, with zoomed-in view on the right side for clearer demonstration. Scale bar: 100$\mu m$. All the data volumes were downloaded from the published article[9].

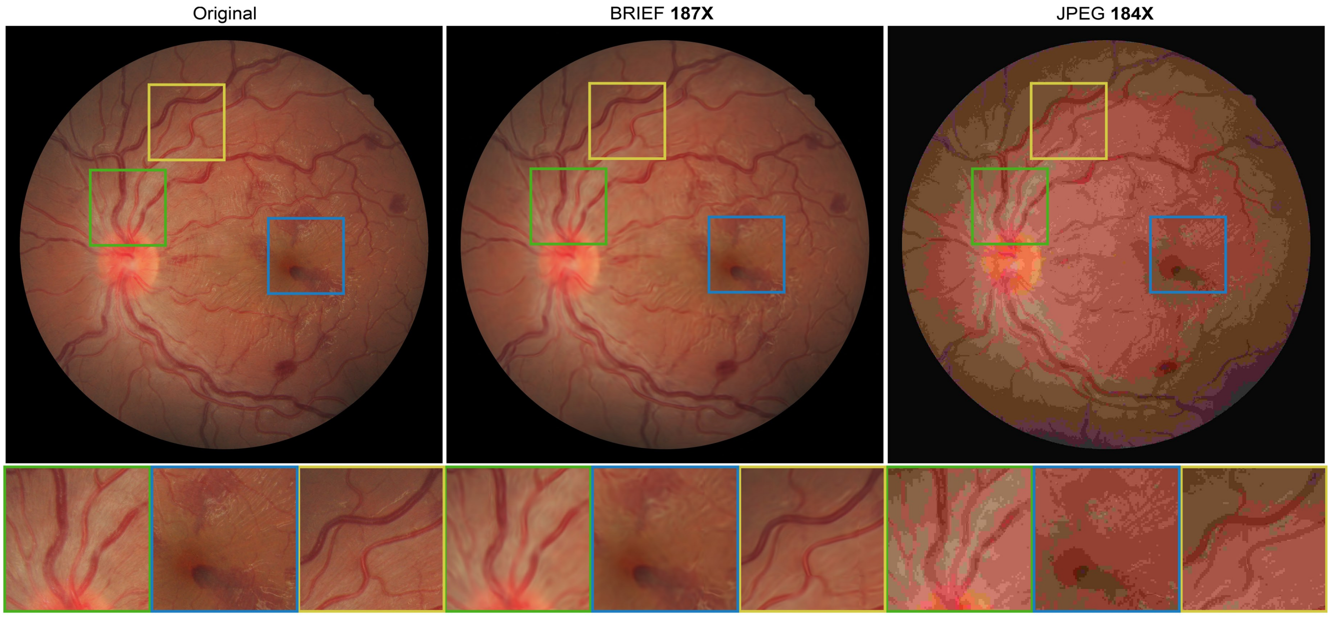

**Supplementary Figure 8. The comparison between BRIEF and JPEG on fundus image.** Top: the visual comparison of the original data and decompressed versions by BRIEF and JPEG for fundus image at around 187× compression ratio. Bottom: the close-up views on ROI are labeled at the top. BRIEF clearly preserves the continuous blood vessels and avoids color distortion. The 2D images were released from the published article[10].

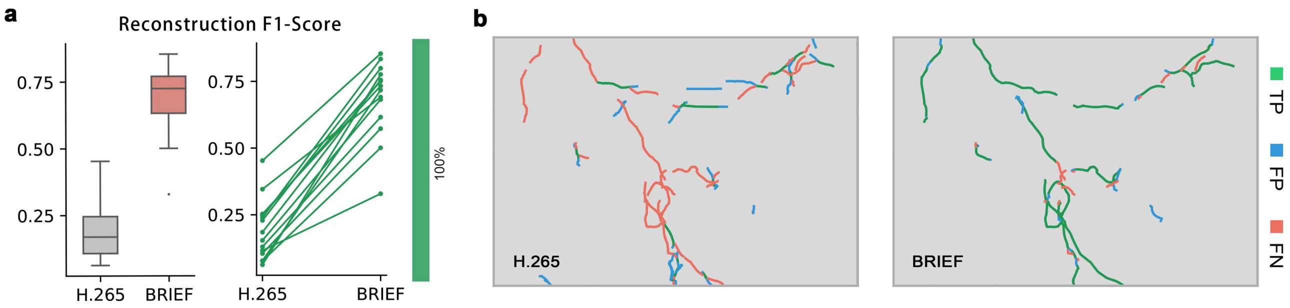

**Supplementary Figure 9. The comparison between BRIEF and H.265 on reconstruction of axonal fibers. a,** The mean and standard variance of Reconstruction F1-score on the traced 3D morphological structure after compression by H.265 and BRIEF at around 420× compression ratio, and the performance difference over 14 samples, in which the BRIEF performs better and worse are highlighted with green and red lines respectively. **b,** The error map of traced 3D axonal fibers on the decompressed data. TP (True Positive): the correctly reconstructed structures; FP (False Positive): the incorrectly hallucinated structures; FN (False Negative): the missing details.

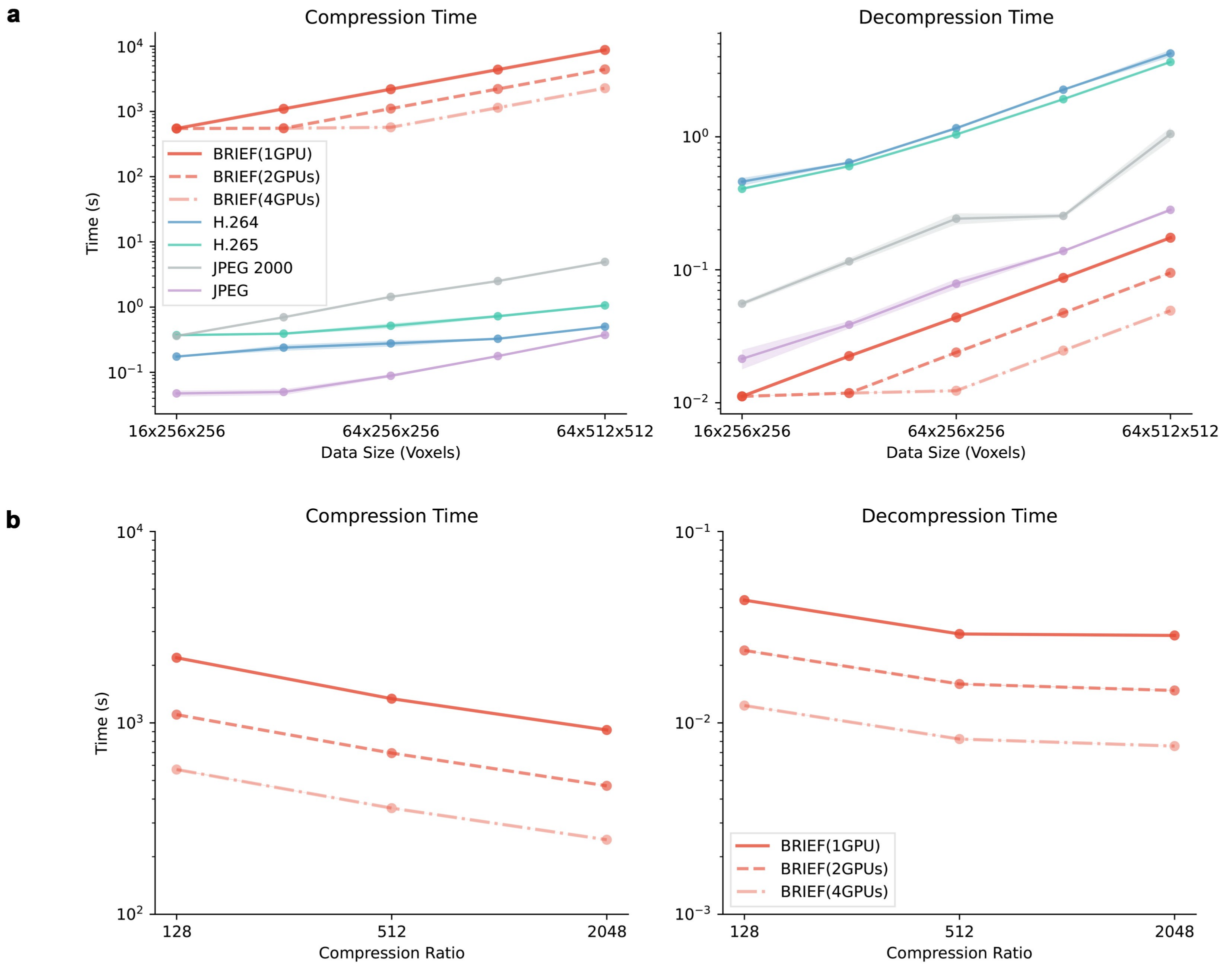

**Supplementary Figure 10. Time complexity analysis for BRIEF. a**, The comparison of BRIEF and state-of-the-art compressors on speed of compression and decompression under around $128\times$ compression ratio and five data size ($16 \times 256 \times 256$, $32 \times 256 \times 256$, $64 \times 256 \times 256$, $64 \times 256 \times 512$, $64 \times 512 \times 512$) for 5 neuronal data volumes. The execution of BRIEF is conducted on GPUs, while the other codecs are CPU-based. The speed of BRIEF with three computing power configurations (1 GPU, 2 GPUs, and 4 GPUs) is also shown. **b**, The speed of compression and decompression of BRIEF under three compression ratios ($128\times$, $512\times$, $2048\times$) with three computing power configurations (1 GPU, 2 GPUs, and 4 GPUs), which adopts the $16 \times 256 \times 256$ neuronal data volumes used in the **a.**

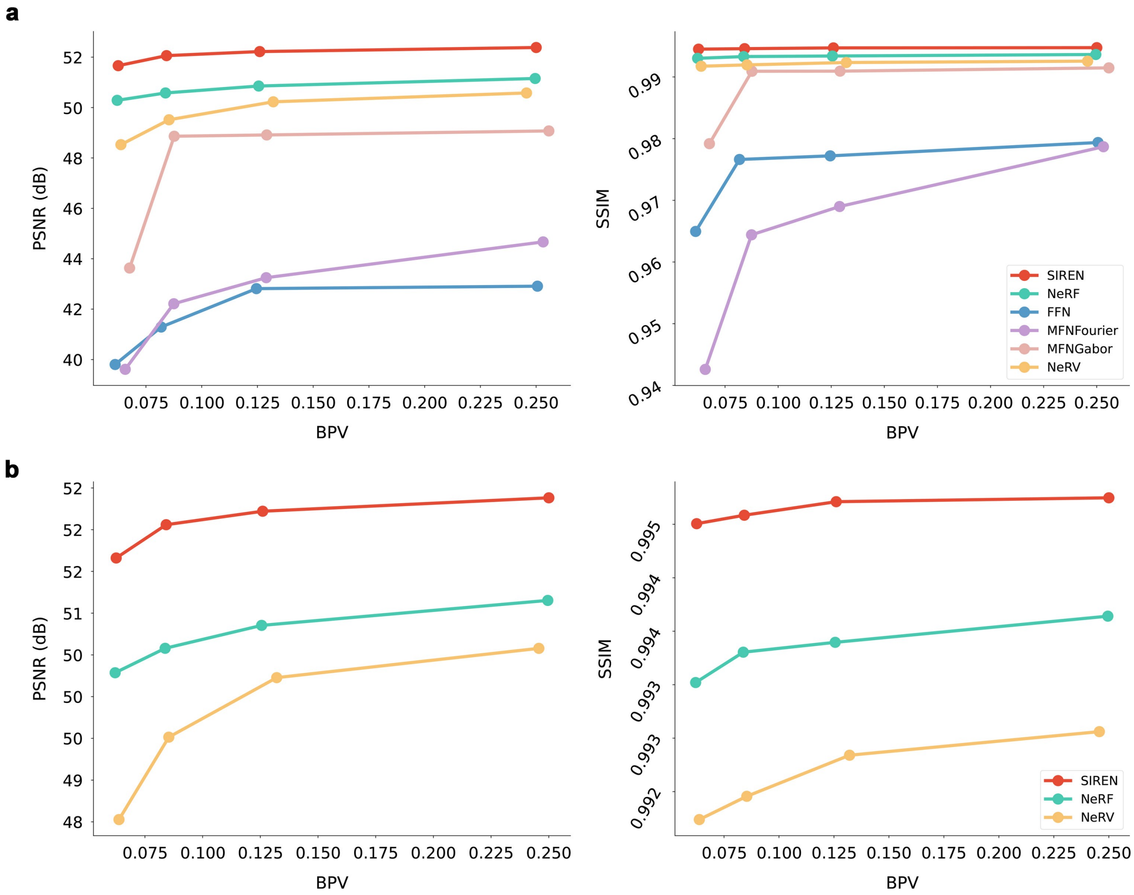

**Supplementary Figure 11. Performance comparison between SIREN and other MLP architectures. a**, The performance of SIREN and other MLP architectures on organic CT data volumes at varying compression ratios. The comparison architectures include NeRF[11], MFNFourier[12], MFNGabor[12], and NeRV[13]. **b**, Performance comparison between the top three architectures (SIREN, NeRF, and NeRV).

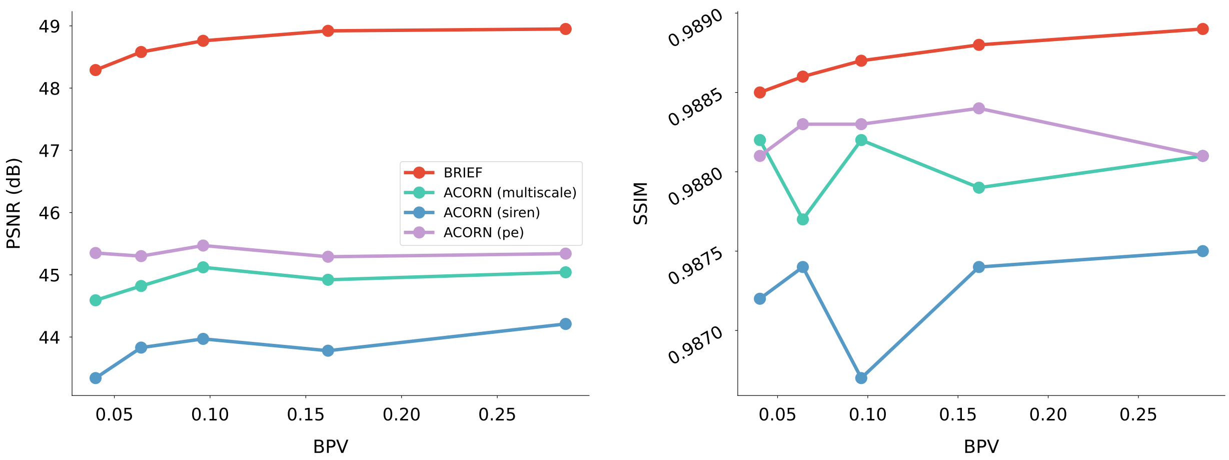

**Supplementary Figure 12. Performance comparison between BRIEF and ACORN.** We cropped the organic CT data into $64 \times 512 \times 512$-pixel blocks, which is the maximum data size viable for running the ACORN algorithm on our GPU RTX 3090 (24GB) and conducted comparative experiments at five distinct compression ratios. To ensure a comprehensive comparison and mitigate the influence of variations in neural network architectures, we contrasted not only the algorithm as originally described in ACORN (multiscale) but also two other neural network structures mentioned in their paper (pe and siren).

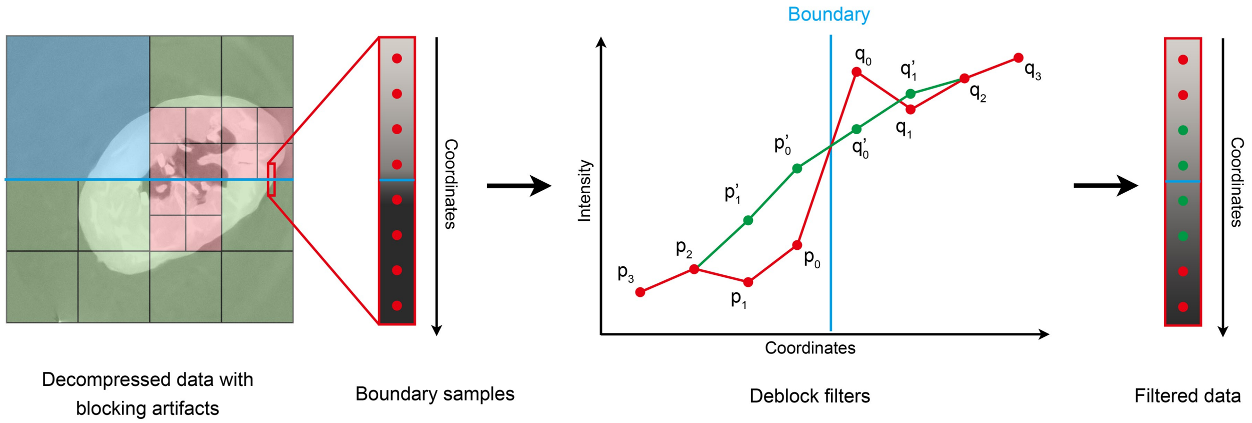

**Supplementary Figure 13. Adaptive deblocking filter.** The decompressed data has obvious block artifacts, and there are vertical and horizontal boundaries between blocks. Take the blue horizontal boundary line in the left figure as an example. First, We take four points from both sides of the boundary and get a total of eight points $p_3$, $p_2$, $p_1$, $p_0$, $q_0$, $q_1$, $q_2$, $q_3$, which are shown in red. Then judge whether to filter these samples according to their values. If the result shows that the boundary does have blocking artifacts, we filter these sampling points and obtain the filtered data $p_1'$, $p_0'$, $q_0'$, $q_1'$, which are shown in green in the figure.

#### 6 Supplementary Tables

**Table 1.** **Implementation details of comparison algorithms.**

| Methods | Implementation Details | Preprocess | Computing Device |
|---|---|---|---|
| JPEG | cv2.imwrite, cv2.IMWRITE_JPEG_QUALITY set to 10-70 | Flattened to 2D image | CPU |
| JPEG 2000 | glymur.Jp2k[1], colorspace='gray', cratios=compress_ratio | Flattened to 2D image | CPU |
| H.264 | ffmpeg -y -f rawvideo -s frame_size -pix_fmt gray16 -r 30 -I - -an -c:v libx264 -b bitrate save_path | Converted to byte stream | CPU |
| H.265 | ffmpeg -y -f rawvideo -s frame_size -pix_fmt gray16 -r 30 -I - -an -c:v libx265 -b bitrate save_path | Converted to byte stream | CPU |
| DVC | Pytorch Implementation[2], trained with $\lambda = 16, 256, 4096, 64000, 1024000$ | I Frames compressed with JPEG (Quality=30) | GPU |
| SGA | Tensorflow Implementation[3], bb_sga configuration, num_filters=192 and 256, trained with $\lambda = 0.001, 0.0025, 0.005, 0.01, 0.02, 0.04, 0.08$ | Flattened to 2D image slices of size 1024×1024 | GPU |

[1] https://pypi.org/project/Glymur/
[2] https://github.com/binzzheng/DVC-PyTorch
[3] https://github.com/mandt-lab/improving-inference-for-neural-image-compression

**Table 2.** **The comparison between BRIEF and state-of-the-art compressors on reconstruction of axonal fibers.** The average and standard variance of Reconstruction F1-score on the traced 3D morphological structure after compression by BRIEF , H.264, H.265, JPEG, and JPEG 2000 at around 420× compression ratio, and the performance difference over 14 samples.

| Methods | Recon. F1 Score (Avg.) | Recon. F1 Score (Std.) |
|---|---|---|
| BRIEF (ours) | 0.617 | 0.202 |
| H.264 | 0.252 | 0.180 |
| H.265 | 0.191 | 0.112 |
| JPEG | 0.200 | 0.174 |
| JPEG 2000 | 0.139 | 0.158 |

**Table 3.** **BRIEF's speed of compression and decompression on two typical biomedical datasets.**

| Data name (compression ratio) | Size (voxels) | Compression time (hours) | Decompression time (minutes) |
|---|---|---|---|
| Whole brain neurons (2244×) | $28965 \times 20005 \times 12000$ | 1794 | 42 |
| HiP-CT (128×) | $3147 \times 1811 \times 1811$ | 23.8 | 0.6 |

**Table 4.** **The comparison of BRIEF and state-of-the-art compressors on speed of compression and decompression.** The execution of BRIEF, DVC, and SGA is conducted on GPUs, while the other codecs are CPU-based. Here we show the mean of compression time and decompression time for each method on 10 neuron data, which were cropped to the same size ($64 \times 512 \times 512$ voxels), under around 512× compression ratio. JPEG was excluded due to the lower compression ratio.

| Methods | Compression (seconds) | Decompression (milliseconds) |
|---|---|---|
| BRIEF (ours) | 1113 | 216.4 |
| H.264 | 0.900 | 448.6 |
| H.265 | 2.414 | 501.1 |
| DVC | 38.99 | 295.6 |
| SGA | 3231 | 279.3 |
